# Evolutionarily divergent mTOR remodels the translatome to drive rapid wound closure and regeneration

**DOI:** 10.1101/2021.10.28.465024

**Authors:** Olena Zhulyn, Hannah D. Rosenblatt, Leila Shokat, Shizhong Dai, Duygu Kuzuoglu-Öztürk, Zijian Zhang, Davide Ruggero, Kevan M. Shokat, Maria Barna

## Abstract

An outstanding mystery in biology is why some species, such as the axolotl, can scarlessly heal and regenerate tissues while most mammals cannot. Here, we demonstrate that rapid activation of protein synthesis is a unique, and previously uncharacterized, feature of the injury response critical for limb regeneration in the axolotl (*A. mexicanum)*. By applying polysome sequencing, we identify hundreds of transcripts, including antioxidants and ribosome components, which do not change in their overall mRNA abundance but are selectively activated at the level of translation from pre-existing mRNAs in response to injury. In contrast, we show that protein synthesis is not activated in response to digit amputation in the non-regenerative mouse. We further identify the mTORC1 pathway as a key upstream signal that mediates this regenerative translation response in the axolotl. Inhibition of this pathway is sufficient to suppress translation and axolotl regeneration. Surprisingly, although mTOR is highly evolutionarily conserved, we discover unappreciated expansions in mTOR protein sequence among urodele amphibians. By engineering an axolotl mTOR in human cells, we demonstrate that this change creates a hypersensitive kinase that may allow axolotls to maintain this pathway in a highly labile state primed for rapid activation. This may underlie metabolic differences and nutrient sensing between regenerative and non-regenerative species that are key to regeneration. Together, these findings highlight the unanticipated impact of the translatome on orchestrating the early steps of wound healing in highly regenerative species and provide a missing link in our understanding of vertebrate regenerative potential.

## Main

The biggest biomedical challenge of this century is the restoration of diseased or damaged organs and tissues. Urodele amphibians, such as newts and salamanders, display the highest regenerative capacity among vertebrates. In contrast, most mammals have limited regenerative potential. The axolotl, in particular, is famous for its extraordinary ability to rapidly regenerate organs, including limbs, hearts, and brains, throughout its lifespan^1^. The mechanisms underlying this diversity in regenerative potential across the animal kingdom remain poorly understood. Our current understanding of regeneration is largely based on the transcriptome^2–5^ and not on direct assessment of proteins that are ultimately required for repair and tissue regeneration. This is in part due to technical limitations - until recently, we lacked the tools to study translation of mRNA into protein at the same scale and resolution as transcription^6–8^. Furthermore, at 32 Gb, axolotls have one of the largest genomes sequenced to date and their genes are on average 25x longer than those of humans^9–11^. The *de novo* transcription and processing of these expanded transcripts may be too slow to guide the rapid responses to amputation, including wound closure, which occur within hours of injury. Therefore, a better understanding of what molecular processes influence the ultimate gene expression program of rapid wound healing is needed and is highly relevant to treating injury, organ failure, and disease.

Here we integrate emerging technologies with experiments in regenerative and non-regenerative species to discover an exquisite genome-wide post-transcriptional gene expression program guiding wound closure and regeneration. These findings pinpoint rapid ‘on-demand’ translational remodeling of distinct gene networks involved in translational control, growth, and metabolic adaptation, which may be particularly important for injury repair and promoting regeneration. We further identify <u>m</u>echanistic target <u>o</u>f <u>R</u>apamycin (mTOR) as a critical regulator, of this translational program, with an essential function in tissue regeneration. Moreover, our data reveal the surprising evolution of the sequence of mTOR, within regenerative species which creates a ‘hypersensitive’ kinase that extends its range of function and sensitivity. This uncovers the unexpected impact of highly conserved, key signaling pathways to be remodeled to foster differential regenerative potential across animal kingdoms with potential implications for our broader understanding of health and disease.

### Amputation of the axolotl limb triggers an increase in protein synthesis before the proliferative stage

The differences between regenerative and non-regenerative species arise at their initial cellular response to a traumatic injury. While species like the axolotl experience rapid migration of epithelial cells to scarlessly seal the wound within a day of amputation (Fig. 1a; Extended Data Fig. 1), non-regenerative species take days to close a wound site, often with extensive scarring^1, 12–14^. Scarless wound closure in turn paves the way for a signaling-competent wound epithelium which stimulates the formation of a blastema – a stem-cell like progenitor pool that develops into a new limb (Fig. 1a)^1^. Failure of regeneration may be viewed as failure to create a regeneration-competent cellular milieu in response to injury^15^. To examine how protein synthesis changes in response to injury in the axolotl, we bisected the forearms of wildtype axolotls and harvested a 2 mm thick tissue slice from the wound site at 0 hpa (hours post-amputation) and 24 hpa. Time-matched tissue was pooled and subjected to sucrose gradient fractionation to separate mRNAs based on their ribosome occupancy, providing a snapshot of translation in the sample (Fig. 1b-c). Gradient traces from tissue harvested at 0 hpa had pronounced monosome (1 ribosome per transcript) peaks and a flat “heavy polysome” (3+ ribosomes per transcript) region indicating relatively low basal translation (Fig. 1b, d). Surprisingly, tissues harvested at 24 hpa were characterized by low monosome peaks and the emergence of “heavy polysome” peaks indicating increased protein synthesis (Fig. 1c-d). This response is particularly notable as there are no changes in proliferation before 2 days post-amputation (dpa)^16–18^, suggesting that increased protein synthesis, at this time point, is not simply associated with new cells and tissues being produced in the course of regeneration (Fig. 2a).

**Figure 1:**
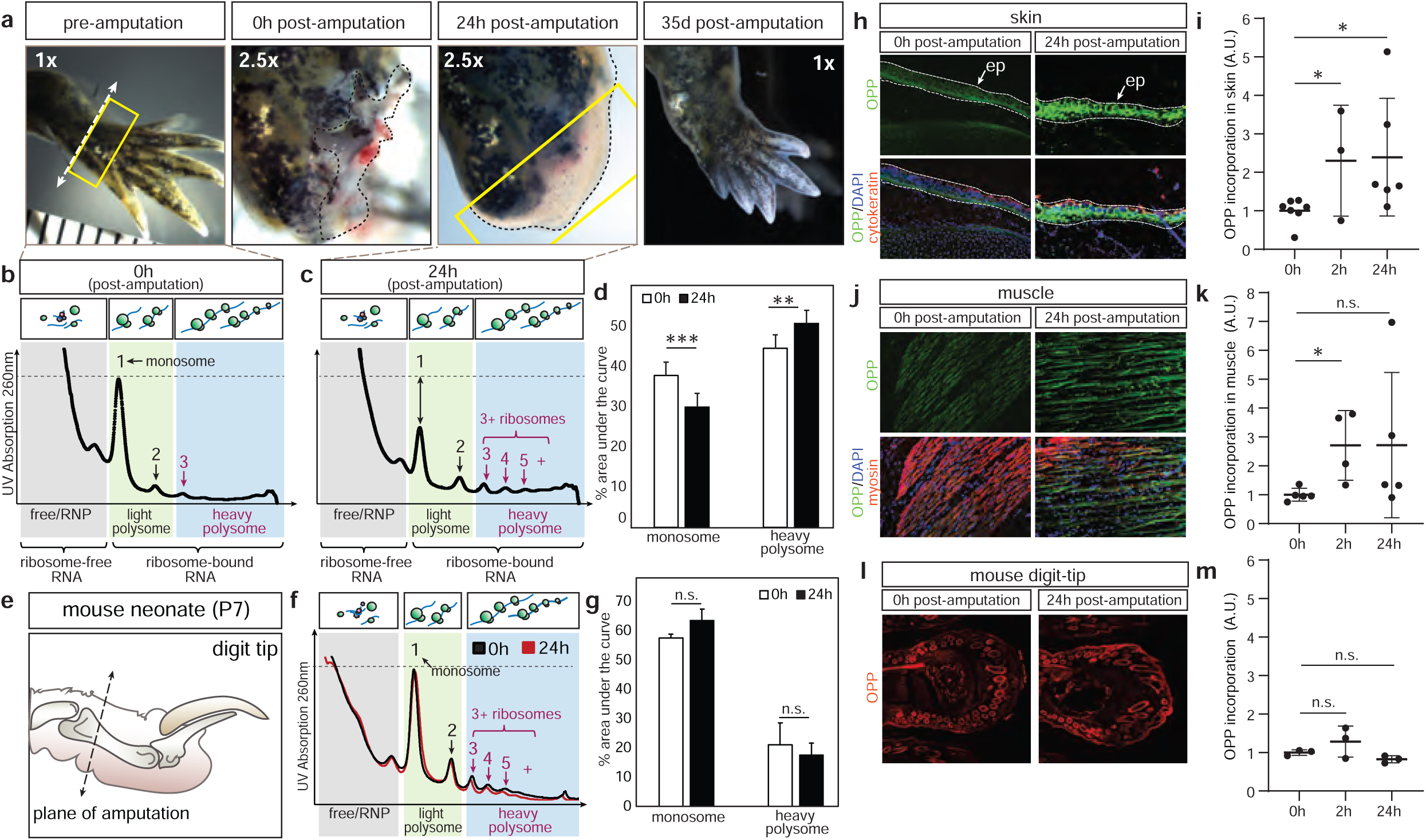
Activation of protein synthesis distinguishes responses to injury in axolotls and mice. **a**, Axolotl forelimbs were amputated at mid-forearm level (dashed line) regenerate in the course of a month. Box shows region of tissue harvested at the plane of amputation. **b**, Sucrose gradient fractionation of tissue harvested at 0 hpa. Dashed line indicates height of monosome peak. Arrows highlight the location of peaks corresponding to 2+ ribosomes. **c**, Arrows highlight drop in monosome peak and increase in heavy polysome peaks after amputation. **d**, Area under the curve quantitation of sucrose gradient traces at 0 hpa and 24 hpa, n=5 per time point. **e**, Schematic of digit tip amputation in the neonatal mouse. Plane of amputation through the 2^nd^ phalanx shown with dashed line. **f**, Sucrose gradient fractionation of mouse digit tips harvest at 0 hpa and 24 hpa. **g**, Quantitation of the gradient trace, n=3 per time point. **h**, OPP incorporation in the skin proximal to the plane of amputation in the axolotl. Dashed line indicates the epithelium (ep). OPP signal indicated by green, cytokeratin staining in red, nuclei in blue. **i**, Quantitation of OPP signal in the epithelium. **j**, OPP incorporation into axolotl muscle near the wound site at 0 hpa and 24 hpa. OPP shown in green, heavy chain myosin (myosin) in red, nuclei in blue. **k**, Quantitation of OPP incorporation in muscle. **l**, OPP incorporation (red) in mouse digit tips at 0 hpa and 24 hpa. **m**, Quantitation of OPP incorporation in mouse digit tips. Statistical significance in **d**, **g**, **i-m** assessed with Student’s t-test. * is p < 0.05, ** is p < 0.01, *** is p < 0.005, n.s. is p > 0.05 and not considered significant.

**Figure 2.**
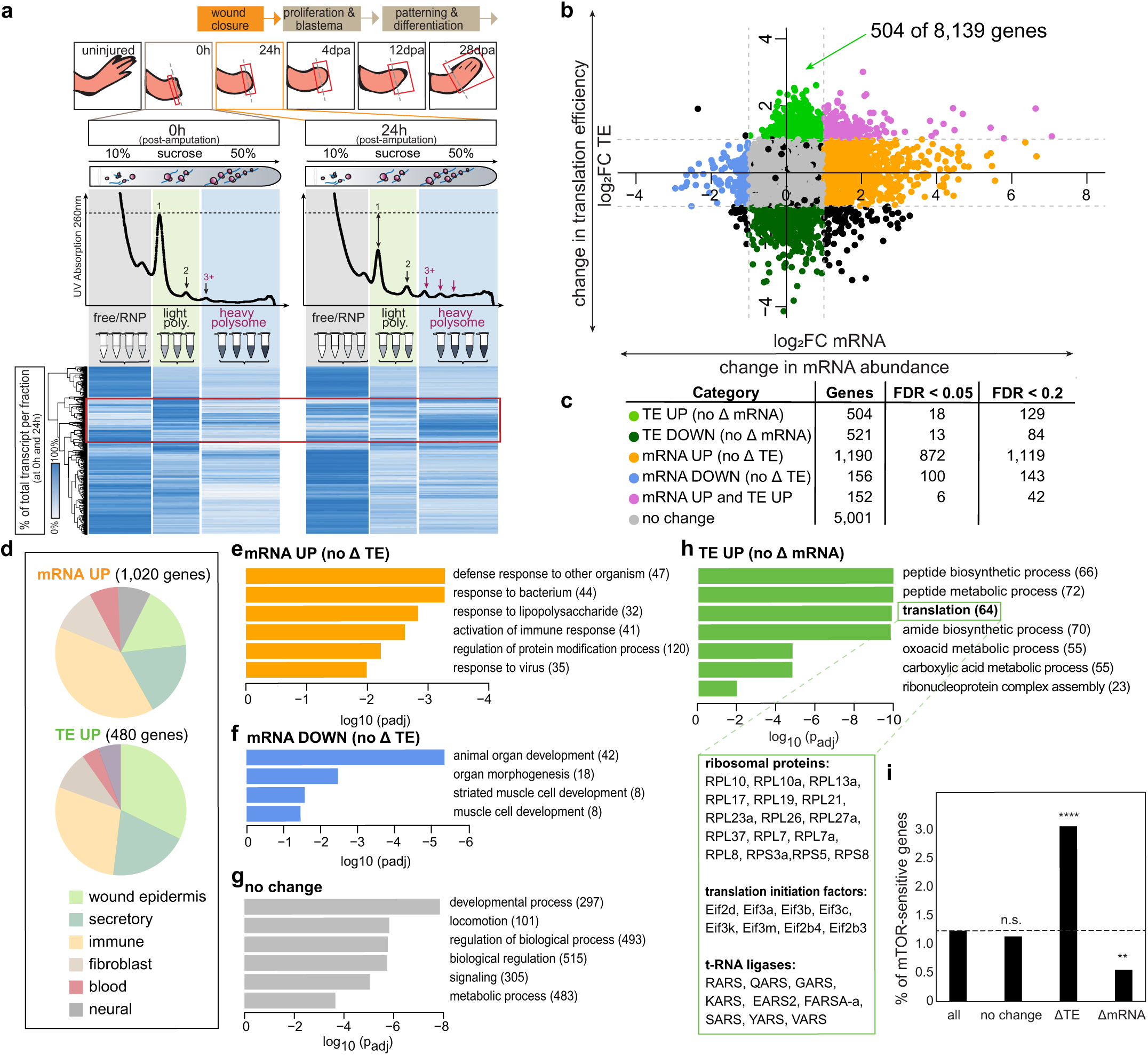
Regenerative translation targets the dormant transcriptome for rapid translational remodeling. **a**, Schematic depicts the polysome sequencing pipeline. Tissues were harvested at 0 hpa and 24 hpa of the “wound closure” phase and pooled (n=2 pools per time-point, 5 tissues per pool). A small quantity of lysate was set aside as the input sample for standard RNA-Seq analysis. The remaining lysate was subjected to sucrose gradient fractionation, fractions were pooled into the “RNP/free”, “light polysome” and “heavy polysome” sets as depicted by grey, green, and blue boxes, respectively, and extracted RNA was subjected to RNA-Seq. Heat map shows average reads of a given transcript as a percentage of total reads in the free/RNP, light and heavy polysome at a given time-point. Red box highlights transcript shifts across fractions in response to amputation. **b**, On the x-axis, scatter plot depicts (log_2_) fold-change (FC) in total mRNA abundance in input samples at 0h and 24h post-amputation. The y-axis depicts (log_2_) FC in translation efficiency (TE). Grey dashed lines indicate two-fold change on the log_2_ scale. Grey dots indicate transcripts with no change in the input or fraction enrichment over time; green dots indicate transcripts with a >2-fold increase in TE but no change in mRNA abundance; dark green dots indicate transcripts with a >2-fold decrease in TE and no change in mRNA abundance; orange dots indicate transcripts that show an increase in transcription but no change in TE, blue dots are samples that decrease in transcription but show no change in TE and pink dots show a concomitant increase in both TE and transcription. **c**, Summary of transcripts in each category that show at least a 2-fold increase or decrease in transcription or TE and the number of transcripts that meet false discovery rate (FDR) cut-offs of 0.05 and 0.2, respectively. **d**, Overlay of our data with previously published single-cell RNA-Seq analysis^21^ suggests that mRNAs predominantly activated at the transcriptional level (orange dots in b) are characteristic of immune cells while those activated at the level of translation (green dots in b) are more typical of the wound epidermis and secretory cells of the skin. **e**, Gene Ontology (GO) enrichment analysis of transcript subsets defined in (b) reveals enrichment of immunity-associated biological processes in the transcriptionally upregulated category. **f**, enrichment of cell-type specific differentiation genes in the transcriptionally downregulated category, **g**, enrichment of developmental processes and signaling genes among mRNAs that do not change in our data set, **h**, enrichment of translation and metabolic processes in the translationally activated gene category. Green box shows a selection of ribosomal proteins (RPs), translation initiation factors, and t-RNA ligases that are part of the GO “translation” category enriched in our data set. **i**, Distribution of orthologues of established TOP/PRTE-domain containing mTOR target genes in our data set shows significant enrichment within translationally regulated gene set based on binomial test. The ** indicates p < 0.01, **** indicates p < 0.00001, n.s. indicates p > 0.05 and is not deemed statistically significant.

To better understand whether increased translation is specific to axolotls or a general response to amputation across vertebrates, we made use of the well-established mouse digit-tip amputation model^19^, which allowed us to assess how tissues respond to severe non-lethal injury. We performed proximal amputations of digit 2 in mouse pups at postnatal day 7 (P7) (Fig. 1e) and assessed translation in pools of mouse digit tip tissue at 0 hpa and 24 hpa by sucrose gradient fractionation. At 0 hpa, the sucrose gradient trace exhibited prominent monosome and heavy polysome peaks suggestive of moderate translation. Notably, unlike the axolotl, the sucrose gradient traces of mouse digit tip tissue at 24 hpa were indistinguishable from the 0 h time point (Fig. 1f-g). These findings reveal a striking difference in protein synthesis at the site of amputation between regenerative versus non-regenerative species.

To visualize which tissues responded to amputation by increasing translation, we injected white axolotls with O-propargyl-puromycin (OPP), an alkyne analogue of the drug puromycin, which incorporates into nascent proteins and can be detected on tissue sections by means of a “click” reaction with an azide-fluorophore conjugate^20^. We observed moderate OPP incorporation throughout the limb prior to amputation (0 h). However, at 2 hpa, we observed a dramatic increase in OPP incorporation in muscle and connective tissue of the amputated limb. This increase was particularly pronounced in the skin near the wound site (labelled with cytokeratin), where it remained significant at 24 hpa (Fig. 1h-i). In muscle (labelled with myosin heavy chain), the trend was maintained but was not significant at this time point (Fig. 1 j-k). These findings indicate that injury leads to increased protein synthesis in the axolotl. In contrast, *in vivo* OPP incorporation studies revealed that wildtype mice do not exhibit any overt differences in protein synthesis at 0 h, 2 h or 24 h post-amputation (Fig. 1l-m). These findings show that activation of protein synthesis distinguishes responses to injury in axolotls and mice.

### Polysome sequencing reveals rapid ‘on demand’ remodeling of gene expression at the translation level upon injury

We next sought to determine whether upregulations in protein synthesis upon axolotl limb amputation was associated with changes in the translation of specific transcripts. We therefore characterized remodeling of gene expression at the level of translational control by optimizing tissue harvest, lysis, sucrose gradient fractionation and analysis to develop an axolotl-specific polysome sequencing pipeline in which transcripts associated with the “free/ribonucleoprotein (RNP)”, “light” and “heavy” polysome fractions were fractionated and sequenced from tissues harvested at 0 and 24 hpa and compared to transcripts from total lysate (“input”) (Fig. 2a). We identified 8,139 mRNAs with detectable reads across samples. Consistent with our observation of increased protein synthesis, 17.3% of transcripts showed a greater than two-fold increase in heavy polysome association, indicating increased translation (Extended Data Fig. 2a). For half of these transcripts, greater polysome association was associated with the overall increase in their transcript abundance after amputation (Extended Data Fig. 2b). To identify transcripts that were selectively translated in response to amputation independent of their transcription status, we calculated the change in translational efficiency (TE - the enrichment of a given transcript in the heavy polysome compared to the free and light fractions combined, calculated as TE = ((heavy24h - heavy0h) - ((RNP24h + light24h) - (RNP0h + light0h))) and expressed on a log_2_ scale) of each mRNA. We identified 504 transcripts that showed a two-fold increase in their TE, but no change in their transcription (calculated as input24h – input0h and expressed on a log_2_ scale), between 0 h and 24 h post-amputation (green dots in Fig. 2b-c; Extended Data Fig. 2b). These transcripts are particularly interesting because they represent pre-existing mRNAs that are recruited to the ribosome in response to injury. A relative increase in TE is predicted by a drop in enrichment in the free/RNP fraction but not overall transcript abundance (Extended Data Fig. 2c-d). This suggests that increased translation of existing mRNAs occurs in response to amputation. Moreover, because they do not change significantly in transcript abundance, these transcripts are overlooked by traditional transcriptome analyses.

When we overlaid our data with a single-cell RNA-Seq data set^21^, we observed that more than 50% of cell-type annotations associated with translationally activated (“TE UP”, green dots in Fig. 2b-c) mRNAs are associated with epidermal cells (including cells of the intermediate and basal wound epidermis shown in green and small secretory cells and Leydig cells shown in blue in Fig. 2d). These data are consistent with elevated OPP incorporation observed in the skin near the wound site (Fig. 1h). Conversely, less than one third of transcriptionally activated (“mRNA UP”, orange dots in Fig. 2b-c) transcripts are expressed in the skin, while more than one third are derived from immune cells (including T cells, B cells, neutrophils, phagocytosing neutrophils and recruited macrophages) (Fig. 2d). Consistent with this, Gene Ontology (GO) enrichment analysis of transcriptionally activated mRNAs revealed that GO terms associated with immune responses; for example, “defense response to other organism”, “activation of immune response” and “lymphocyte migration”, are highly enriched within the “mRNA UP” gene set (Fig. 2b orange; Fig. 2e; Extended Data Table 2). This GO signature likely reflects both migration of immune cell types to the wound and activation of inflammation markers at the wound site^3, 18, 22–24^. Whereas 5,001 mRNAs that show no changes in mRNA abundance or TE after amputation were highly enriched in regulators of “signaling” and “development”, suggesting that these transcripts are not deployed until wound closure is completed (Fig. 2b grey; Fig. 2g; Extended Data Table 2; Extended Data Fig. 2e)^3, 18^. Of note, 521 genes in our data set belonged to the “TE DOWN” gene set and showed a marked decrease in translation efficiency without a change in mRNA abundance. This gene set did not show any significant enrichment of GO terms with the caveat that >66% of the mRNAs were not annotated (“TE DOWN” dark green dots in Fig. 2b; Extended Data Table 2). Broadly, these findings highlight that translation and transcription target distinct biological processes in response to amputation. This dichotomy may reflect differences in time-scales upon which these processes operate such that cells may utilize translation of pre-existing transcripts for rapid, on-demand production of critical proteins and transcription to drive longer-term processes, such as immune activation.

Of the translationally upregulated transcripts, the top 20 included several key regulators of cellular redox state, including thioredoxin (TXN) and peroxiredoxin (PRDX), which play important roles in disulfide bond reduction and negative regulation of reactive oxygen species (ROS)^25^, and SELENBP1a (methanethiol oxidase), which generates hydrogen peroxide in the process of oxidizing methanethiol^26^ (Fig. 2b-c green, Extended Data Table 1). While excessive levels of ROS are detrimental to cellular homeostasis, previous studies show that at a certain threshold ROS act as important activators of signaling pathways. The translational control of ROS regulators is interesting as their activation at sites of injury is an important facet of regeneration in a variety of species, including zebrafish, frogs, planaria, and axolotls^27–33^. Selective translation of these proteins suggests that induction and a delicate balance of ROS during wound healing and regeneration is a part of a heretofore unexplored and tightly regulated post-transcriptional regulatory process. Our data also show that a key regeneration regulator, anterior gradient protein 2 (AGR2), the first known nerve-derived factor established to be a driver of blastema induction and regeneration in newts, is selectively translated as early as 24 h post-amputation, indicating that it may also play a role in early wound healing (Extended Data Table 1)^2, 21, 34^.

To gain insight into the biological processes involved in wound healing, we also conducted GO enrichment analysis on our set of translationally activated pre-existing mRNAs. To our surprise, these transcripts were highly enriched in GO terms associated with protein synthesis, such as “translation”, “peptide biosynthetic process” and “ribonucleotide complex assembly” and included key regulators of this process (Fig. 2b-c green; Fig. 2 h; Extended Data Table 2). For example, the “translation” GO term included multiple ribosomal proteins, translation initiation factors and t-RNA ligases (Fig. 2 h inset, Extended Data Table 1). This hinted at a possible mechanism, downstream of a distinct signal, that coordinates the translation of key mRNAs encoding components of the protein synthesis machinery after injury. Indeed, it is well-established that translation of many ribosomal proteins and translation initiation factors is directly regulated by complex 1 of the mTOR pathway through pyrimidine-rich translational element (PRTE) and 5′ terminal oligopyrimidine tract (TOP) motifs encoded in the 5’ leader sequences of their mRNAs^35^. We identified 101 axolotl orthologues of established mammalian mTORC1 targets (Extended Data Table 3)^36, 37^ in our data set and observed that they are disproportionately enriched within the set of mRNAs that show a change in TE after amputation, compared to those that change only in their mRNA abundance or show no change at all (Fig. 2i). Therefore, activation of mTOR signaling may be a central driver of translational remodeling upon amputation.

### Injury is associated with rapid activation of mTOR signaling in the axolotl

mTOR activation stimulates quiescent stem cells^38^ and has been observed during the proliferative phases of zebrafish, axolotl and planarian regeneration^17, 39, 40^. However, mTOR activation during the early wound healing phase has not been characterized in axolotl. The mTOR pathway integrates an array of environmental cues such as nutrient abundance, oxygen levels, growth factors and stress, to regulate cellular growth and metabolism. The serine/threonine kinase mTOR forms the catalytic core of two distinct complexes, mTORC1 and mTORC2. While, mTORC2 is primarily involved in regulation of proliferation and cytoskeletal rearrangements, mTORC1 is a master regulator of protein synthesis^41–43^. Activation of mTORC1 triggers phosphorylation of two sets of translational regulators, S6Ks (p70 S6 kinase 1 and 2) and 4E-BPs (eukaryotic initiation factor 4E-binding proteins 1, 2 and 3)^41, 43, 44^. Phosphorylation of S6K1 activates its kinase activity and it, in turn, phosphorylates ribosomal protein S6 (RPS6) at five highly conserved sites (Ser235, 236, 240, 244 and 247), which are used as markers of mTORC1 activation^45^. Western blot analysis of axolotl tissue harvested from the wound site between 0 h and 24 h after amputation demonstrated a linear increase in phosphorylation of both Ser235/236 and Ser240/244 residues between 2 h and 24 h post-amputation (Fig. 3a-b). Of note, while Ser235/236 may be phosphorylated in both an mTORC1-dependent and independent manner, the phosphorylation of the other residues depends exclusively on S6K1 and its homologue S6K2^45, 46^, demonstrating that amputation of the axolotl limb robustly activates the mTORC1 pathway.

**Figure 3:**
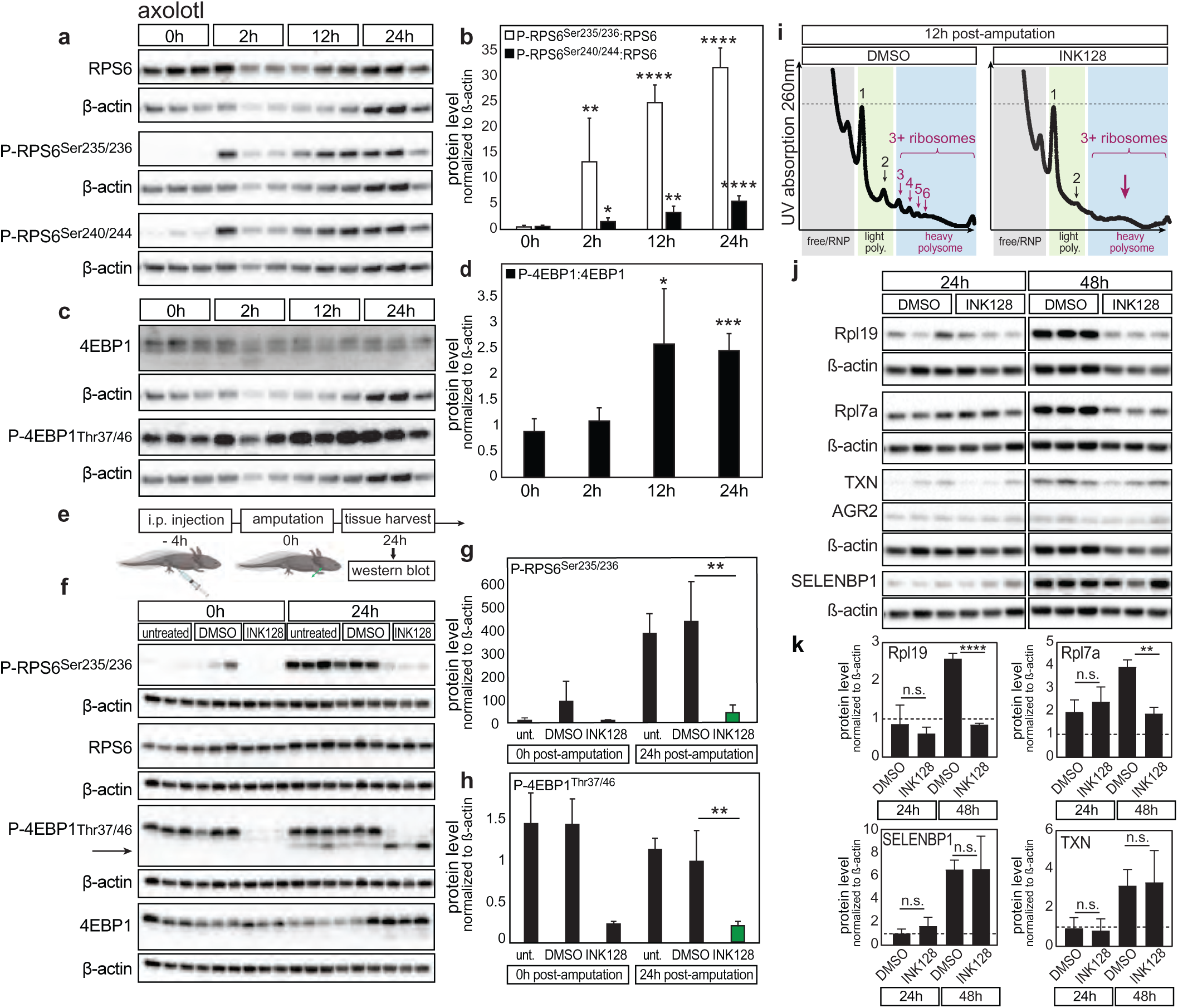
Injury is associated with rapid activation of mTOR signaling in the axolotl. **a**, Limb tissue lysates harvested at indicated time-points after amputation. Each lane contains tissue from a distinct animal assessed by western blot for changes in mTOR activity using antibodies against total RPS6, phosphorylated (P)-RPS6^Ser235/236^, P-RPS6^Ser240/244^, and ß-actin. **b**, Quantitation of phosphorylated RPS6 normalized to ß-actin relative to total RPS6 normalized to own ß-actin. **c**, Western blot of P-4EBP1^Thr37/46^, total 4EBP1, and ß-actin. **d**, Quantitation of phosphorylated 4EBP1 normalized to ß-actin relative to total 4EBP1 normalized to own ß-actin. Note that RPS6 and 4EBP1 were sequentially blotted on same membrane therefore share the same ß-actin; P-RPS6^Ser235/236^and P- of P-4EBP1^Thr37/46^ were sequentially blotted on the same membrane therefore share the same ß-actin. For b and d, significance was assessed with Student’s t-test comparing 0 hpa to 2 hpa, 12 hpa, or 24 hpa for each target protein and p-value is indicated above the bar for the given time-point. The * is p < 0.05, ** is p < 0.01, *** is p < 0.001, **** is p < 0.0001. **e**, Schematic illustrates timing of intraperitoneal (i.p.) injection of drug and amputation in the axolotl. **f**, Western blot analysis of tissues harvested from axolotls treated with INK128, DMSO carrier or untreated. Each lane contains tissue from distinct animal. P- RPS6^Ser235/236^ and 4EBP1 were sequentially blotted on the same membrane and share ß-actin, RPS6 and P-4EBP1^Thr37/46^ were sequentially blotted on the same membrane and share ß-actin. Arrow highlights P-4EBP2 band. **g**, Quantitation of P-RPS6^Ser235/236^ normalized to ß-actin. “unt.” refers to untreated animals that did not undergo i.p. injection. **h**, Quantitation shows levels of P- 4EBP1^Thr37/46^ normalized to ß-actin. In g-h, significance was assessed with Student’s t-test comparing protein level in DMSO to INK128 treated animals at 24 hpa, ** is p < 0.01. **i**, Representative sucrose gradient traces from (n=3) independent experiments assessing translation in DMSO or INK128-treated animals at 12 hpa. Numbered arrows indicate location of monosome (1), disome (2), and heavy polysome (3+) peaks. Large arrow highlights flattened heavy polysome (poly.) region upon INK128-treatment. **j**, Western blot analysis of translationally regulated genes, in tissues from n=3 distinct animals per condition, shows decreased protein levels of Rpl19 and Rpl7a in INK128-treated animals at 48 hpa. No significant decrease in TXN, AGR2, and SELENBP1 levels upon INK128 treatment at either 24 hpa or 48 hpa. **k**, Quantitation of protein levels normalized to ß-actin. Dashed line indicates protein level in DMSO controls at 0 hpa. Significance assessed with Student’s t-test comparing indicated pairs of data. The ** is p < 0.01, **** is p < 0.0001, n.s. is p > 0.05 and deemed not significant.

While phosphorylation of RPS6 serves as a sensitive marker of mTORC1 pathway activation, its role in translation remains unclear^45^. Conversely, phosphorylation of 4E-BPs by activated mTORC1 plays a critical role in regulating protein synthesis^47–50^. In its native state, unphosphorylated 4E-BP1 sequesters the cap-binding protein eIF4E, thereby inhibiting translation. Phosphorylation of 4E-BP1 releases eIF4E to form the translation initiation complex and stimulates 5’ cap-dependent translation^41, 47, 48^. Axolotl tissues express high levels of both total and phosphorylated 4E-BP1 at steady state. However, we observe a significant increase in 4E-BP1 phosphorylation between 12 h and 24 h post-amputation, coincident with the onset of epithelial migration (Fig. 3c-d). These data suggest that the spike in protein synthesis observed during wound closure may be directly regulated by mTORC1 activation.

To test this hypothesis we treated axolotls with INK128 (MLN0128), a potent ATP-site specific inhibitor of mTOR^36^. Four hours after administering the drug, we amputated the left forelimb and assessed mTORC1 activation by western blot at 24 hpa (Fig. 3e). Compared to controls (untreated or DMSO carrier control), axolotls treated with INK128 showed robust inhibition of RPS6 and 4E-BP1 phosphorylation within 4 h of drug treatment and failed to activate the mTORC1 pathway in response to limb amputation (Fig. 3f-h). Sucrose gradient fractionation of tissue harvested from the wound site revealed a profound decrease in translation in INK128-treated animals compared to DMSO-treated controls at 12 hpa (Fig. 3i). These data indicate that the increase in protein synthesis observed after amputation is partly dependent on mTOR activity. To assess if selective translation of pre-existing transcripts in our polysome sequencing data was dependent on mTORC1, we performed western blot analysis on tissues harvested from INK128-treated animals at 0 h, 24 h and 48 h post-amputation. We observed a significant decrease in the protein levels of known mTORC1-targets, Rpl19 and Rpl7a^36, 37^, at 48 h post-amputation in drug-treated animals compared to controls. In contrast, we did not observe a significant change in the levels of TXN, AGR2 or SELENBP1 in response to INK128 (Fig. 3j-k). This suggests that selective translation encompasses both mTORC1-dependent and independent mechanisms. Although, certain transcripts may rely on alternative mechanisms to maintain a high level of translation in the presence of the inhibitor.

### mTOR is required for wound healing and regeneration

To determine the role of mTORC1 signaling in axolotl wound closure we examined the effect of INK128 treatment on wound healing after limb amputation. Treatment with INK128 was associated with profound deficits in wound closure in 87.5% of drug-treated axolotls at 24 hpa. In contrast, 100% of controls (both untreated and those treated with DMSO carrier control) exhibited complete wound closure (Fig. 4a-b). Live imaging of wound closure revealed that treatment with INK128 impaired epidermal cell migration between 30 min and 90 min after amputation (Fig. 4c; Extended Data Video 1 (control); Extended Data Video 2 (INK128-treated)). These findings demonstrate that mTORC1 activation is critical for rapid wound closure observed within 24 h of amputation.

**Figure 4:**
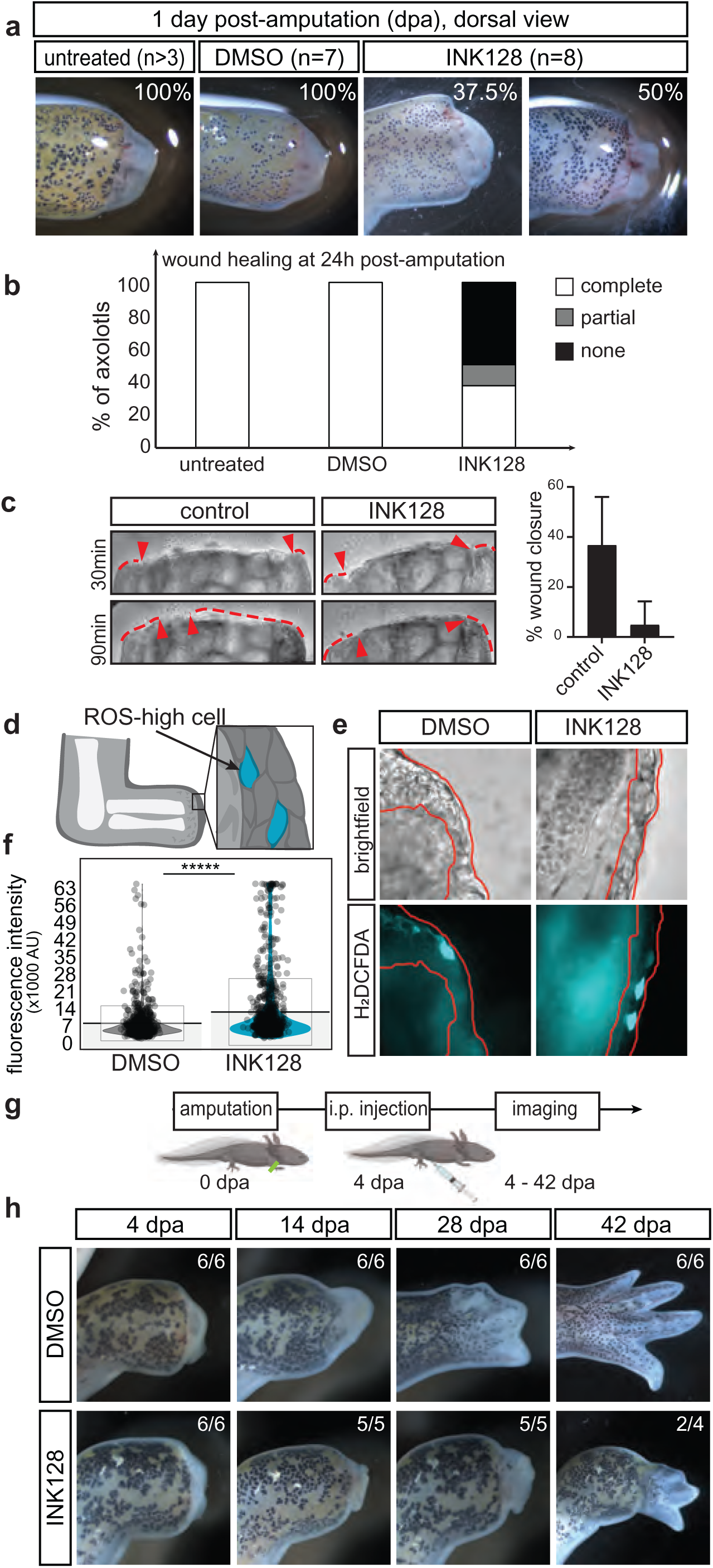
mTOR activation promotes rapid wound closure and regeneration. **a**, Representative images of wound site morphology in untreated (n>3), DMSO (n=7) or INK128 (n=8) treated animals at 1 day post-amputation with % of animals achieving full wound closure shown in panel. **b**, Quantitation of wound closure phenotypes upon INK128 treatment. “complete” indicates full closure of the wound site with multiple layers of cells apparent; “partial” indicates that wound site is closed with a single layer of cells at the apex; “none” indicates failure of cell migration over the plane of amputation and evidence of protruding bone. **c**, Representative captures of cell migration over the wound site at 30 min and 90 min post-amputation in control (representative video in Extended Data Video 1) and INK128-treated animals (Extended Data Video 2). Quantitation shows percentage of wound closure at 1 hpa. Red arrows indicate leading edge of skin on either side of the wound site. Dashed line highlights skin adjacent to the wound site. **d,** Schematic shows distribution of reactive oxygen species (ROS) high cells (blue) and low cells (grey) at the wound site. **e**, Representative images of ROS levels in cells, labeled with H_2_DCFDA (blue), at the wound site at 36 hpa, after treatment with DMSO or INK128. The epithelial layer is outlined in red. **f**, Quantitation of H_2_DCFDA fluorescence intensity in the epithelial layer outlined in red. Points represent intensity of individual labelled cells at the wound site. Significance assessed with Student’s t-test, **** is p < 0.0001. **g**, Schematic illustrates experimental set-up to assess role of mTOR beyond wound closure. Axolotls underwent amputation at 0 dpa and all (12/12) showed complete wound closure at 4 dpa. Animals were treated with either INK128 or DMSO carrier at 4, 6, 8 and 10 dpa. **h**, Representative images of axolotls treated with DMSO or INK128 at 4dpa as described in g and imaged at 4, 14, 28 and 42 dpa.

Our polysome sequencing, detected translational activation of peroxiredoxin, an important antioxidant, within 24 hpa. Previous studies showed that sustained accumulation of ROS is critical for tissue regeneration ^29–31, 33^. In contrast, high levels are toxic, therefore cells must carefully orchestrate a balance of ROS production ^51^. To address whether inhibition of translation impacts the ROS balance *in vivo*, we used 2’,7’-dichlorodihydrofluorescein diacetate (H_2_DCFDA) to detect ROS levels in INK128 or DMSO-treated limbs at 36 hpa (Fig. 4d-f). We observed a significant increase in the intensity of H_2_DCFDA after drug treatment which suggested that inhibition of mTORC1 resulted in an increase in ROS at the wound site (Fig. 4e-f). Together, these results indicate that mTOR activation influences key aspects of the early tissue regenerative response such as wound closure and appropriate accumulation of ROS levels.

In addition, we observed that even transient effects on mTORC1 signaling had long-term effects on regeneration. For example, the animals which responded to a single dose of INK128 with partial wound closure at 24 hpa, developed further tissue regression between 4 and 7 dpa and delayed induction of a blastema compared to controls (Fig. 4a; Extended Data Fig. 3). This implied that sustained mTORC1 signaling is required after wound closure as suggested by detection of phosphorylated RPS6 during the proliferative phase (3-14 dpa)^23^. To address this hypothesis, and to rule out the possibility that regeneration is delayed due to the regression of the wound epithelium which is required for blastema induction^1^, we allowed axolotl limbs to undergo complete wound closure before treating the animals with INK128 at 4, 6, 8 and 10 dpa (Fig. 4g-h). However, treatment with INK128 after wound closure was completed did not prevent tissue regression at 14 dpa. Furthermore, at 28 dpa, all INK128-treated animals remained arrested at the blastema stage while the controls had well-developed limb buds with distinct digit rays (Fig. 4h). This indicated that sustained mTORC1 signaling is required not only to drive initial wound closure immediately after injury, but also to maintain tissue integrity and sustain blastema formation during the proliferative stage. Notably, the drug-treated animals were able to re-initiate regeneration and form digits between 42 and 56 dpa likely due to subsequent re-activation of the mTOR pathway.

### Evolution of hypersensitive mTOR kinase underlies divergent nutrient sensing and tissue regenerative response

To determine if amputation-induced activation of mTORC1 is unique to regenerative axolotls or if it is a general response observed in vertebrate wound healing, we examined the status of mTORC1 signaling in digit tips of neonatal mice using the toe-clipping assay described in Fig. 1. Western blot analysis of digit tip tissue at 0 hpa, 2 hpa, and 24 hpa revealed that while mTORC1 signaling is basally active, it does not show a further increase in activation in response to injury (Fig. 5a-b; Extended Data Fig. 4a; Extended Data Fig. 5) suggesting that mTORC1 activation, and concomitant increase in protein synthesis, are a response that is specific to regenerative axolotls.

**Figure 5:**
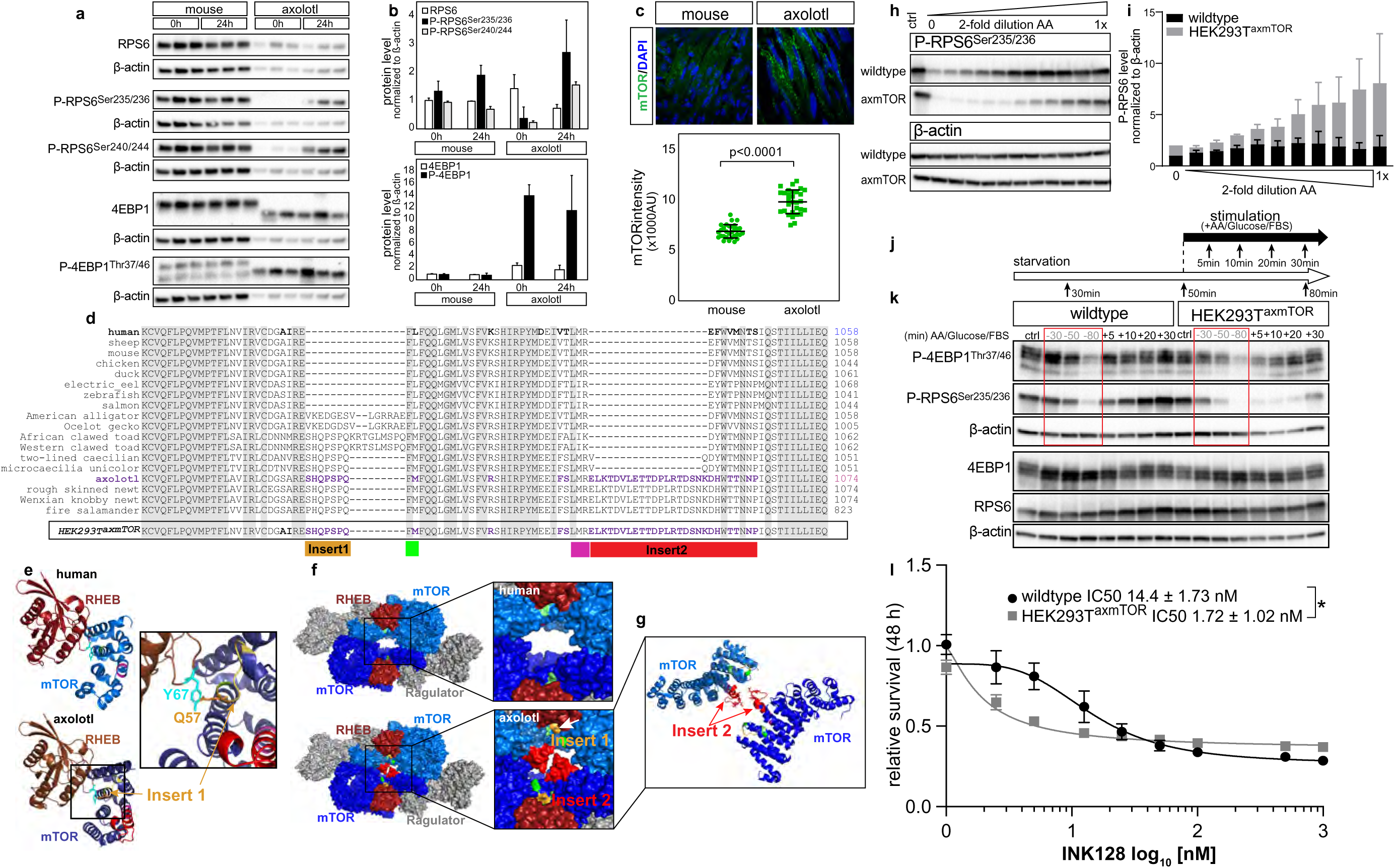
Urodele-specific insertions give rise to hypersensitive mTOR kinase. **a**, Western blot analysis shows high basal mTOR activity but no significant increase in phosphorylated RPS6 or 4EBP1 in mouse digit tips at 24 hpa, in contrast to pronounced mTOR activation in the axolotl upon amputation. **b**, Quantitation of mTOR target phosphorylation in the axolotl and mouse. Protein level normalized to ß-actin. **c**, Diffuse, granular immunofluorescent staining for mTOR (green) observed in mouse muscle at steady state. In contrast, in axolotl muscle cells, mTOR is coalesced in pronounced puncta. The average mTOR intensity is quantified from 10 sections across 3 individual animals per species. **d**, Subset of a multiple sequence analysis across metazoan species illustrates the location of urodele specific insertions in the axolotl (magenta). Box highlights the sequence of the homozygous chimeric HEK 293T^axmTOR^ cell line. Insert 1 shown in yellow, insert 2 in red, green and magenta boxes show adjacent conserved residues. **e**, Structure of human RHEB-bound mTOR (PDB 6BCU) and predicted structure of axolotl RHEB-bound mTOR near the switch II region. Insert 1 is shown in yellow with Q57 residue labelled. RHEB Y67 is labelled in blue. Inset highlights the new interaction interface. **f**, Surface structure of the human Rheb-bound mTORC1 dimer (6BCU) illustrates insert 2 (red) filling the cavity formed by mTOR dimer (blue). **g**, Cartoon view highlights the relative orientation predicted for insert 2 in M-HEAT region of antiparallel mTOR proteins within the dimer. In e-g, insert 1 is yellow, insert 2 is red, subunits of mTOR dimer are royal blue and blue respectively, RHEB is burgundy, Ragulator is grey, conserved residues adjacent to insert 1 are green, conserved residues adjacent to insert 2 are pink. **h**, Representative western blot of amino acid (AA) titration experiment emphasizes increased sensitivity of P-RPS6^Ser235/236^ phosphorylation in HEK 293T^axmTOR^ to amino acid concentration. **i**, Quantitation of P-RPS6^Ser235/236^ level. **j**, Schematic of time-course experiment. Cells undergo combined amino acid, glucose and serum starvation for a total of 80min. “Starved” samples are harvested at 30 min, 50 min and 80 min (lanes labelled -30, -50, and -80 respectively). After 50min of starvation (dashed line), cells are stimulated with a 1x concentration of amino acids, glucose and serum and samples and are harvested at 5, 10, 20 or 30 min (lanes labelled with +). **k**, Representative western blot (n=2 distinct cell line clones) analysis of starvation/stimulation time-course described in j. Control cells at steady state are labelled “ctrl". Red box highlights the profound drop in P-4EBP1^Thr37/46^ and P-RPS6^Ser235/236^ phosphorylation observed in HEK 293T^axmTOR^ cells upon starvation. Mutant cells exhibit a more graded response to stimulation with respect to P-4EBP1^Thr37/46^ and reduced phosphorylation of P-RPS6^Ser235/236^ suggesting altered substrate preference. **l**, survival assay illustrates that axolotl insertions confer hypersensitivity on human mTOR kinase.

To understand where the difference between axolotls and mice arises, we examined the mTORC1 pathway itself. It has been established that in the presence of nutrients, mTORC1 is recruited to the lysosomal membrane via the Rag-Ragulator complex where it is activated by RHEB and phosphorylates its substrates (Extended Data Fig. 4b)^52–54^. The components of this cellular machinery are highly conserved with axolotl proteins exhibiting an average of ∼88% amino acid identity to their human and mouse orthologues (mTOR, RAPTOR, MLST8, LAMPTOR1-5, RAGA, RAGC, RAGD and RHEB) (Extended Data Table 4). However, we noticed an interesting feature when we examined the intracellular distribution of endogenous mTOR within tissues. In the mouse limb, at homeostasis, the mTOR staining appears diffuse and cytoplasmic (as shown in muscle in Fig. 5c). In contrast, axolotl mTOR localizes to lysosomes across multiple cell types and its localization diminished upon INK128 treatment (Fig. 5c, Extended Data Fig. 4c-d)^52, 54^. This suggests that, in contrast to mammalian cells, a pool of axolotl mTOR remains constitutively localized to the lysosome potentially priming this pathway for more rapid activation.

Although mTOR is highly conserved, to understand if there are specific features exclusive to axolotl mTOR, we performed a multiple sequence alignment across 50 metazoans, predominantly vertebrates, including 8 species of newts and salamanders (Extended Data Figure 6, Supplementary Information). This analysis revealed that while axolotl mTOR is highly conserved (∼84 % amino acid identity compared to both its human and mouse orthologues) (Extended Data Table 4), it contains two intriguing insertions embedded within otherwise highly conserved and functionally important M-HEAT regions encoded by exon 20 and 21 of mTOR (Fig. 5d). Insert 1 is an ∼8 residue sequence within the M-HEAT region of axolotl mTOR which is critical for RHEB-mediated activation^59–62^. Structure modelling based on the available human mTORC1 structures (PDB 6BCU) suggests that Gln57 within insert 1 may make a new direct contact with RHEB at Y67 of its Switch II domain that is critical for activating mTORC1 (Fig. 5e). Notably, while this insertion is highly conserved across all three extant orders of amphibians (frogs, caecilians and salamanders), it is expanded by 6 residues in frogs. In addition, reptiles appear to encode a distinct insertion at this locus. The insertion is otherwise absent across metazoans and the mammalian sequence appears to be ancestral as it is also found in invertebrate species. In contrast, insert 2 is a ∼20 residue insertion, within the M-HEAT domain, which is found exclusively in newts and salamanders (Fig. 5d-e). This insertion extends the M-HEAT and N-HEAT interface, predicted to play a role in protein-protein interactions (Fig. 4f-g)^55^. To assess the impact of these insertions, we used CRISPR/Cas9-mediated gene editing to engineer a chimeric human-axolotl mTOR by incorporating the two contiguous axolotl insertions in frame into the human mTOR genomic locus (axmTOR) to generate homozygous HEK293T^axmTOR^ cell lines (Fig. 5d boxed sequence; Extended Data Fig. 7a-c). Subsequent nutrient deprivation experiments revealed that in HEK293T^axmTOR^ cells, mTOR localization to lysosomes is less sensitive to starvation than in the wildtype HEK293T and that a pool of mTOR persists at lysosomes of HEK293T^axmTOR^ cells, similar to what we observe in the axolotl (Extended Data Fig. 7d-e; Fig. 5c).

Based on our structural modelling of axolotl mTOR, which suggests that it may have altered interactions with RHEB, and our observation that the chimeric mTOR persists at the lysosome, we hypothesized that axolotl insertions may alter the enzymatic activity of mTOR kinase. It is established that mTOR is activated upon nutrient and amino acid stimulation^56–58^. Notably, nutrient signaling plays an important role in tissue regeneration^59–64^ and it has recently been shown that nutrient cues alone are sufficient to induce regeneration in a variety of species^59^. By carrying out amino acid titration experiments, we show that HEK293T^axmTOR^ cells are far more effective than wildtype HEK293T at sensing the decrease in nutrient levels as well as responding to subtle changes in amino acid concentration by incrementally regulating the phosphorylation of 4EBP1 and RPS6 (Fig. 5h-i, Extended Data Fig. 7f-g). HEK293T^axmTOR^ cells are also better able to discern differences in duration of exposure to nutrients by modulating phosphorylation of mTOR substrates (Fig. 5j-k; Extended Data Fig. 7h-i). Of note, the HEK293T^axmTOR^ cells exhibit more robust phosphorylation of 4EBP1 than RPS6 across all time-points and concentrations of amino acids. These findings suggest that axolotl-specific insertions alter mTOR’s ability to sense and transduce incremental concentrations of nutrients. Therefore, the salamander-specific mTOR insertions may allow axolotl cells to maintain an increased dynamic range of mTOR activation to more rapidly and robustly sense nutrient and metabolite changes underlying injury and cellular processes associated with regeneration.

Lastly, a viability assay illustrates that the chimeric mTOR kinase is hypersensitive to INK128 treatment and exhibits a significantly lower IC50 than its human counterpart (Fig. 5l). Despite being a ‘hypersensitive kinase’, the chimeric mTOR has the remarkable ability, upon amino acid titration, to not exceed the upper limit of activation of the wildtype. This finding suggests that it is not functioning at the level of an oncogenic mTOR kinase. Intriguingly, this notion is consistent with the axolotl’s capacity to regenerate but not elicit uncontrolled growth, a feature that can be coupled with its extraordinary resistance to oncogenic transformation^65, 66^. Therefore, the functional differences in upstream regulation of protein synthesis control, through mTOR sequence differences between species, may underlie the remarkable ability to repurpose a classic ‘stress-response’ signal to a ‘growth and regeneration’ signal.

## Discussion

In this study, we examined the translational landscape of wound closure in a regenerative species, revealing for the first time the hidden repertoire of hundreds of pre-existing transcripts that are rapidly recruited to ribosomes for translation in response to amputation. The identification of mTORC1 targets within our data set not only provides mechanistic insight into how some of these transcripts may be regulated but also paves the way to expanding the known repertoire of mTORC1-target genes outside the oncogenic context. We further show a critical role of mTOR signaling in wound healing and regeneration. In contrast to the mammalian kinase which readily reaches maximum activity, axolotl mTOR integrates a wide range of nutrient concentrations to more incrementally dial up or dial down its activity. Predictive modelling of axolotl mTOR onto the human structure positions Q57 of insert 1 in direct contact with Y67 and close to D65 of the RHEB switch II domain. Previous studies showed that Y67 is one of just four RHEB residues that are essential for mTOR kinase activity^67^. In contrast, D65 is required to promote the GTPase activity of RHEB which is intrinsically low compared to that of other small GTPases, thereby favoring a high activation state for this protein^68^. Thus, it is plausible that the positioning of Q57 and insert 1 reorients the switch II domain to stimulate GTPase activity thereby conferring greater sensitivity to the RHEB-mTOR complex, but not exceeding maximum activity.

Analysis of mTOR sequences from a variety of lizards, turtles, birds, crocodiles and snakes reveals that most species encode a highly conserved alternative at the same position as the amphibian insert 1 (VKEDGDSVLGRRAE). The distribution of these insertions across amphibians and reptiles suggests the possibility that insert 1 arose in cold blooded species as a means to titrate back the high activity of the ancestral one which remains highly conserved across warm blooded species. One possible explanation of this evolutionary divergence is that the intrinsically high basal activity of the ancestral mTOR kinase was not compatible with the environmental conditions and metabolic needs of cold-blooded species. Previous studies have demonstrated that there is a direct trade-off between the high metabolic activity required to sustain warm blooded species and regenerative potential^69^. The reduction of regenerative potential may reflect a lack of resources to sustain tissue replacement under conditions of rapid metabolism. Additionally, the comparatively low sensitivity and narrow dynamic range of mammalian mTOR may mean that it may not discern or respond to activating cues associated with injury because these are masked out by the high basal activity of this kinase in warm-blooded species. It has recently been shown that nutrient cues alone, including insulin and leucine, are sufficient to induce regeneration in a variety of species, including non-regenerative mice^59^. As leucine in particular is a powerful stimulator of mTOR, this suggests that increased sensitivity of this pathway in the axolotl may have a rapid and direct beneficial effect in coupling nutrient sensing with gene expression at the post-transcriptional level to initiate and sustain regeneration^59, 69–73^.

Just as our understanding of lysosome localization extended our knowledge of mTOR function, in this work we illustrate that although mTOR is both critical and highly conserved, it shows remarkable evolutionary malleability that extends its range of function and sensitivity. Understanding how evolutionary tweaks have altered the capacity of amphibians to balance metabolic and regenerative demands may also shed light on regulatory mechanisms employed by these species, particularly the axolotl, to develop a natural cancer resistance^74^. Historically, the lack of broad regenerative ability across the animal kingdom has been attributed to a belief that tissue regeneration is inextricably linked to cancer. The axolotl provides a curious counterpoint to this paradigm as this animal, and related urodele species, possess both prodigious regenerative abilities and a surprising resistance to cancer^65, 66^. Our identification of urodele-specific changes in functional regions of mTOR, a kinase often deregulated in many tumors, provides an ideal starting point to better understand the mechanistic relationship between regeneration and cancer in the context of mTOR-mediated metabolism and gene expression. These findings further suggest the possibility that engineering an axolotl-like mTOR activity may allow us to promote wound healing, and perhaps even tissue regeneration, in mammals without triggering oncogenic transformation.

## Methods

### Animal experimentation

Animal experiments were approved by Stanford University’s Institutional Animal Care and Use Committee. Axolotls were housed in 40% Holtfreter’s Salt Solution (“tank water”) at room temperature. Axolotls were anesthetized before any procedure involving amputation, injection or live imaging using Tricaine dissolved in 40% Holtfreter’s Salt Solution at the following concentrations titrated based on body size (0.25 g/L for animals 3-8 cm snout to tail tip, 0.5 g/L for 8-12 cm animals, 1 g/L for 12-15 cm animals and 2 g/L for 15-19 cm animals). Experiments were performed at 16-20°C. Axolotls recovered in 0.5 mg/L butorphanol tartrate dissolved in clean tank water for 24 h after the procedure. Animals were euthanized following APLAC approved procedures (axolotls in 5 g/L Tricaine; mice via CO_2_ exposure). Axolotl limb amputations were performed through the left forelimbs at the mid-forearm level. Mouse digit-tip amputations were performed at the level of P2.

### OPP Incorporation and Immunofluorescent Staining

O-Propargyl Puromycin (OPP) was purchased from Medchem Source LLP (25mg, JA-1024), reconstituted in sterile DMSO (100 mM final concentration), aliquoted and stored at -80°C. Prior to injection, OPP aliquot was further diluted in DMSO (Sigma) to a working concentration of 10 mM (4.9553 mg/mL). Animals were anesthetized, weighed and underwent amputation as described. OPP was injected intraperitoneally 1h prior to tissue harvest at 0 h, 2 h, or 24 h post-amputation. Injection aliquots were prepared by combining appropriate volume of 10 mM stock and 0.8x sterile PBS for axolotls (or 1x sterile PBS for mice) adjusted to a pH of 6.5. Next, 0.49553 mg of OPP was administered per 20 g body mass. An injection volume of 100 μL was used. After 1 h, animals were euthanized following APLAC approved protocol. Tissue was harvested and fixed for 1 h at 4°C in 4% paraformaldehyde in PBS. For all washes, 0.8x PBS was used for axolotl tissues and 1x was used for mice. Tissues were washed 4x for 15 min at 4°C. The first wash was in PBT (PBS, 0.1% Tween), subsequent washes were in PBS. Samples were equilibrated in 30% sucrose solution in filtered 0.1 M potassium phosphate buffer at 4°C overnight, and for 1 h in Tissue-Tek O.C.T. (Sakura Finetek, #24583) in molds. Samples were frozen on dry ice. Tissues were sectioned, equilibrated to RT (room temperature) for 1h and rehydrated for 1h in blocking buffer (PBS with 1% goat serum and 0.1% Triton-X) and incubated overnight, at 4°C, with primary antibody against cytokeratin 5/6/18 (Santa Cruz Biotechnology, sc-53262) or myosin heavy chain (DSHB, MF-20) at 1:50 dilution in blocking solution. Slides were washed three times 15 min in blocking buffer and incubated with secondary antibody conjugated with Alexa Fluor 488 or Alexa Fluor 568 (Thermo Fisher, A-11017 or A10037) at 1:500 dilution and DAPI at 1:1000 dilution in blocking buffer for 1h in the dark at RT. Slides were rinsed twice in blocking buffer for 15 min and twice in PBS for 5 min. The Click-iT Alexa Fluor Picolyl Azide Toolkit was reconstituted, and components were combined per manufacturer’s instructions (Thermo Fisher C10641 or C10642). Components C and D were combined in a 1:4 ratio. Slides were rinsed twice in PBS for 5 min and 150 μL of reaction cocktail as added per slide. Slides were incubated for 30 min in a humidified chamber in the dark, at RT. They were then rinsed twice for 5min in PBS and coverslips were mounted with Fluoromount-G Slide Mounting Medium (Southern Biotech, 0100-01). Slides were imaged on a spinning disc confocal microscope and images were acquired with Volocity. Mean fluorescence intensity was measured within a manually defined region of interest (ROI) containing the digit at the plane of amputation (mice), skin (defined by cytokeratin staining) or muscle (defined by MHC) staining. Mean background fluorescence was also measured in an adjacent ROI lacking tissue and subtracted from the sample fluorescence. Mean fluorescence intensity of a given sample is the average of two ROIs normalized to the mean fluorescence intensity at 0 h post-amputation. For analysis in mice n=3 (CD1 at P7) individual animals were used for each time-point. For each time-point, the number of white axolotls (AGSC_101J812) used was n=7 (0h), n=3 (2 h), and n=6 (24 h). Significance was assessed using Student’s t-test.

For immunofluorescent staining shown in Figure 5 and in Extended Data Figures 4c-d, we followed the standard protocol described above but omitted the OPP administration and detection steps, and used anti-rabbit mTOR (Cell Signaling Technology, #2983S) and anti-mouse Rab7 (Santa Cruz Biotech, sc-271608) primary antibodies with Alexa fluor conjugated secondary antibodies at 1:500 dilution as described above. Slides were images on a Zeiss Axio Observer Z1 microscope coupled to the Perkin Elmer UltraVIEW Vox spinning disk confocal microscopy system.

### Sucrose Gradient Fractionation

Axolotl limb amputations were performed as described above. A ∼2 mm piece of tissue was harvested at 0 h and at 24 h post-amputation, immediately frozen by direct immersion in liquid nitrogen and stored at -80°C until use. Samples were prepared by pooling and powderizing tissues in from 2-5 age-matched animals. Tissues were powderized in liquid nitrogen with a mortar and pestle. We prepared n=5 independent pools of tissues for each time-point from 5 wildtype subadults (AGSC_100S) for experiments in Figures 1 and 2. Wildtype juveniles (AGSC_100J812) were used for sucrose gradient fractionation experiments upon DMSO or INK128 treatment shown in Figure 3 where n=3 independent pools of tissue from pairs of animals were used for each treatment. For mouse sucrose gradients, n=3 independent pools of digit 2 tips were used per time-point harvested from CD1 pups at post-natal day 7. Dissociation and lysis of tissues was optimized for the axolotl as follows powderized tissue was resuspended in lysis buffer (100 mM Tris pH7.5, 15 mM MgCl_2_, 150 mM NaCl, 1% Triton X-100, 2 mM DTT, 750U/mL SuperaseIn RNase Inhibitor (Ambion, AM2696), 0.2 mg/mL cycloheximide (Sigma-Aldrich, C7698-1G), 1X Halt Protease and Phosphatase Inhibitor Cocktail (Thermo Fisher, 78442), 25 U/mL Turbo DNase (Thermo Fisher, AM2238), 8% Glycerol, 0.5% DOC, 0.5 mg/mL heparin, 200 U/mL RNaseOUT Recombinant Ribonuclease Inhibitor (Thermo Fisher, 10777019), 400U/mL RNasin (Promega, N2611) at 500 μL per 70 mg of tissue powder. High quantities of RNase inhibitors were used due to allow isolation of intact polysomes which were otherwise degraded by the high endogenous nuclease activity of axolotl tissues. Samples were incubated on ice for 20 min with sporadic vortexing and lysate was pipetted through a p1000 tip and p200 tip and then passed three times through a 16 G sterile needle and four times through an 18G needle to dissociate the tissue. It was then cleared by centrifugation (four times 5 min each at 4°C) at 5000 g. After clearing, the final volume was topped up to 500 μL if needed. For samples to be used for polysome sequencing (see below), 75 μL were removed, combined with 1 mL of Trizol Reagent (Invitrogen, 15596018) and stored at -20°C. The remaining lysate was loaded on 4 mL sucrose gradients (112.5 μL per gradient) and underwent centrifugation for 2.5 h at 4°C at 35,000 rpm on Beckman SW60 rotor. Sucrose gradients were prepared by sequentially layering and freezing 800 μL of 17.5%, 25.6%, 33.8%, 41.9% and 50% sucrose dissolved in 15 mM Tris-HCl pH7.4, 15 mM MgCl_2_, 300 mM NaCl. Gradients were stored at -80°C and thawed overnight at 4°C prior to use. Gradients were fractionated using a density gradient fraction system (Brandel, BR-188). Quantitation was performed by baselining the gradients and calculating the area under the curve for the monosome and heavy polysome (3+ ribosomes) and normalizing it to the total area under the curve under from monosome to the end of the heavy polysome. Significance was assessed by Student’s t-test with values considered significant when p<0.05.

### Library preparation for Polysome Sequencing

Polysome sequencing was performed on two biological replicates which were subjected to independent harvesting, library preparation and sequencing. For each replicate, we prepared an independent pool of tissues for each time-point each containing tissues from 5 wildtype subadults (AGSC_100S) and subjected them to sucrose gradient fractionation as described above. For each time-point lysate was divided across three 4mL sucrose gradients and fractions were collected into tubes containing 100 μL of 10% SDS. Then 500 μL of each fraction was transferred to a tube containing 700 μL acid-phenol, 100 μL water and 100 μL sodium acetate. “Input” samples collected prior to fractionation as described above were combined with 1 mL of Trizol Reagent, 200 μL of chloroform and shaken for 15 s. Input samples were then incubated at RT for 2-3 min and 500 μL of the aqueous phase were transferred to a tube containing acid phenol, water and sodium acetate. After shaking/vortexing briefly, both input and fraction samples were allowed to rest at RT for 5 min and centrifuged at 13,200 rpm for 10 min at 4°C. Then, 400 μL of each fraction sample was combined with isopropanol and GlycoBlue. Samples were gently inverted and precipitated overnight at -80°C. The samples were then centrifuged at 13,200 rpm for 30 min at 4°C. The pellets were washed twice with 75% ethanol, air dried and resuspended in 10 μL (for fractions) or 40 μL (for input) of Ultra Pure DNase/RNase-Free Distilled Water (Invitrogen, 10977-015). For each time-point, the fractions were pooled into three tubes as follows: (1) RNP to 40S, (2) “light” polysome containing the monosome and disome fractions, (3) “heavy” polysome containing all fractions with 3+ ribosomes. The total volume was topped up to 181 μL with Ultra Pure water and 1 μL of the sample was used to check concentration. Then, 1 μL of 1:200 ERCC RNA Spike-In Mix (Invitrogen, 4456740) was added to each fraction pool and concentration was measured again. DNase treatment was performed with TURBO DNase (Ambion, AM2238) for 30 min at 37°C in a total volume of 500 μL. Samples underwent a second acid phenol extraction as described above, were resuspended in 30 μL of Ultra Pure DNase/RNase-Free Distilled Water. The libraries were prepared by the Stanford Genomic Services Center per manufacturer’s instructions using the TruSeq Stranded Total RNA Library Prep with Ribo-Zero Gold Human/Mouse/Rat (Illumina) and sequenced (PEx150) on a HiSeq 4000 (Illumina).

### Analysis of Polysome Sequencing Data

Quality control of raw sequences was performed using FastQC (www.bioinformatics.babraham.ac.uk/projects/fastqc/). Adapters were clipped using Trimmomatic-0.36 with the following settings LEADING:5 TRAILING:40 MINLEN:50^75^. Reads with phred quality score > 33 were kept and FASTQC was repeated again to confirm quality of remaining reads. The axolotl transcriptome^5^ was downloaded from https://portals.broadinstitute.org/axolotlomics/Axolotl.Trinity. CellReports2017.fasta and concatenated to the sequences of ERCC Spike-In mix sequences (Invitrogen, ERCC92.fa) and used to build a transcript to gene reference index with RSEM version 1.2.30^76^. Paired reads were aligned to the index with Bowtie 2 version 2.2.9^77^ and expression was calculated with RSEM. The data set contained 1,388,890 gene models identified in the reference transcriptome. Of these, 473,373 had > 0 reads across all libraries. Normalization of raw read counts in the free/RNP, light and heavy polysome libraries was performed linear scaling to equalize ERCC reads across these libraries to the ERCC reads in the heavy polysome library at 0h. The input at 24h library was ERCC normalized to the input at 0h library. We subset the data to retain transcripts with > 1 cpm across all the input libraries and > 10 reads in total across all RNP/free, light and heavy polysome libraries thereby retaining 8,139 transcripts that had > 0 reads across all libraries and replicates. Differential expression analysis was performed using the edgeR and limma packages^78, 79^. The voom function was used to enable linear modeling of RNA-Seq data with limma^80^. Analysis was performed using the following design matrix (∼0+group+batch) where group is the sample type (i.e. input at 0 h) and batch identifies the specific replicate. The following comparisons were defined in the corresponding contrast matrix to compare the translational efficiency at the 0 h and 24 h time-point TE=((heavy24h-heavy0h)-(RL_24h-RL_0h)) where “heavy” denotes the average expression in the heavy polysome at the given time-point and RL denotes the average expression in the free/RNP (R) and light polysome (L) fractions at a given time-point. This is reported as “Δ TE” in Extended Data Table 1. Change in mRNA abundance in the input at 0 h and 24 h was defined as (input24h-input0h) this is reported in the “Δ mRNA abundance” field of Extended Data Table 1. Additional contrasts defined in the matrix included the change in enrichment in the free/RNP fraction relative to the other two fractions, defined as (RNP24h-RNP0h)-(LH_24h-LH_0h) where RNP refers to the average expression in the RNP/free fraction and LH refers to the average expression in the light and heavy polysome at a given time-point which was used to examine the correlation between TE and enrichment in the free/RNP fraction (log2FC free/RNP) shown in Extended Data Fig. 2c and reported as “Δ RNP/free enrichment” in Extended Data Table 1. This metric was used to determine the Pearson correlation between change in TE and enrichment in the free/RNP fraction compared to the change in mRNA abundance as shown in Extended Data Figure 2c and 2d, respectively. We also calculated the enrichment in the light polysome defined as (light24h-light0h)-(RH_24h-RH_0h) where light refers to the average expression in the light polysome at a given time-point and RH is the sum of average expression in the free/RNP and heavy polysome fractions. This data was used when defining the subset of genes with “no change” in either the input or the free/RNP, light or heavy polysome fractions and depicted as grey dots in Figure 2b. Using the contrast matrices and experimental design described above, linear models were fitted to the data using limma-voom with empirical Bayes method. The p-values were adjusted for multiple-testing using the Benjamini-Hochberg procedure and the adjusted p-values (FDR) are reported for each gene in Extended Data Table 1. Note that in this table, “Gene” refers to the axolotl transcript annotation from the Bryant et al. (2018) assembly, whereas Uniprot, Gene_ID, Protein_Name, Gene_Name, Organism and Gene_Synonyms refer to gene annotations in the closest orthologous gene (for details see Bryant et al. 2018) across a wide-variety of organisms. Note that many genes do not have annotations and therefore have blank space or NA in these respective fields.

In Figure 2b and 2c (and Extended Data Table 1), the green dots highlight the TE UP (no Δ mRNA) category (504 genes) defined as ΔTE > 1 and |Δ mRNA abundance| < 1; dark green dots highlight the TE DOWN (no Δ mRNA) category (521 genes) defined as ΔTE < (-1) & |Δ mRNA abundance| < 1. The mRNA UP and TE UP category highlighted in pink is defined as ΔTE > 1 and ΔmRNA > 1 and contains 152 genes; the mRNA UP (no ΔTE) category highlighted in orange contains 1190 genes and is defined as |ΔTE| < 1 and ΔmRNA > 1. The mRNA DOWN (no ΔTE) category shown in blue is defined as |ΔTE| < 1 and ΔmRNA < (-1) and contains 156 genes; lastly the no change category highlighted in grey contains 5,001 genes and is conservatively defined as all transcripts with |ΔTE| < 1 and |ΔmRNA| < 1 and |ΔRNP/free| < 1 and |Δ light polysome| < 1. Note that due to this stringency, there are some black dots within the grey region that are excluded by this definition.

The change in abundance of a given transcript in the heavy polysome (log_2_ fold change (FC) heavy polysome) is shown on the x-axis in Extended Data Figure 2a-b with the log_10_ adjusted p-value (p_adj_) plotted on the y-axis. Note that the change in the heavy polysome is a less informative metric, than change in TE (as shown in Figure 2b), to describe the translational regulation of a transcript. This is because while 17.3% of transcripts show a significant increase (p_adj_ <0.05) in their abundance in the heavy polysome after amputation (Extended Data Figure 2a), the increase for most is due to a concomitant increase in transcription as illustrated by the distribution of orange and pink dots representing transcripts that increase in their abundance in the input at 24 h post amputation in Extended Data Figure 2b. Whereas analysis of changes in TE clearly identifies transcripts that are upregulated solely at the level of translation (green dots in Extended Data Figure 2b and Figure 2b).

### Gene ontology annotation

Biological function gene ontology annotation was carried out using the Database for Annotation, Visualization, and Integrated Discovery (DAVID, https://david.ncifcrf.gov/home.jsp) which integrates information from a variety of databases such as KEGG, UniGene and Gene Ontology, to provide comprehensive annotation of gene lists^81^. DAVID supports analysis of lists containing genes from a variety of species allowing us to directly use the data associated with orthologue annotations in the Bryant et al. (2018) axolotl transcriptome assembly. We extracted the Uniprot identifiers for orthologues of axolotl genes in each gene subset defined in Figure 2b-c and Extended Data Table 1, for example orange/ΔmRNA UP and green/ΔTE, and compared gene ontology enrichment within each of these sets to a “background” list comprised of Uniprot identifiers of orthologues of genes in the “no change” (grey) category stringently defined as transcripts that do not show change in the input, free/RNP, light or heavy polysome fraction over time. Figure 2e-h show the top 6 biological function categories enriched within each gene set and the full list of significantly enriched functional categories may be found in Extended Data Table 2. Significance was assessed in DAVID using the Benjamini and Hochberg method to determine p-values adjusted for multiple testing using a linear step-up approach^82^. An adjusted p-value < 0.05 was deemed statistically significant and is reported in Extended Data Table 2. Note that although the TE DOWN/no Δ mRNA gene subset (dark green dots in Figure 2b) contains 521 genes, only 182 contained annotations and no significant gene ontology enrichment was observed relative to the background. Further, DAVID annotation and manual search were used to identify Gene Ontology categories associated with cell “signaling” or “development” (i.e. GO:0060173-limb development) and identified 1,995 genes within our data set that belonged to these categories. Next, we examined the distribution of these “signaling” and “development” genes across the ΔTE and ΔmRNA categories defined in Figure 2b. We observed that the majority (>76%) of these genes fell within the “no change” gene cluster (shown in grey in Figure 2b), an additional 14.3% of these genes belonged to the transcriptionally activated gene cluster (shown in orange in Figure 2b) whereas all other categories accounted for fewer than 10% of the remaining genes within the “signaling” and “development” cluster.

### Cell type enrichment analysis

For cell type enrichment analysis, our data was overlaid with a previously published single-cell RNA-Seq analysis of wound healing and regeneration in the axolotl limb and leveraged axolotl gene annotations shared between the two data sets which were mapped to the same transcriptome^21^. We identified 480/504 annotations for the TE UP/no Δ mRNA (green) category and 1,020/1,190 for the mRNA UP (no Δ TE) (orange) category and summarized their distribution across epidermal cell types (including intermediate wound epidermis, basal wound epidermis, Leydig cells and small secretory cells), immune cells (including T cells, early B cells, neutrophils, B cells, recruited macrophages and phagocytosing neutrophils), vascular system cells (including erythrocytes and endothelial cells), neural cells (including dendritic and Schwann cells) as well as fibroblast-like blastema cells.

### mTOR-sensitive orthologue enrichment

To assess distribution of mTOR target genes in our data set, we identified previously described 189 genes shown to contain an-mTOR sensitive TOP or PRTE motif or to exhibit sensitivity to drugs targeting mTORC1 (Extended Data Table 3)^36, 37^. We identified 79 unique axolotl orthologues of these mTOR-sensitive genes within our data set. An additional 11 mammalian mTOR-sensitive genes had two orthologues each in axolotls. The distribution of mTOR-sensitive orthologues across our data set was as follows: 101 mTOR-sensitive orthologues /8139 total genes; 57/5001 in the “no change” data set (grey dots in Figure 2b); 32/504 in the TE UP/no Δ mRNA data set (green in Figure 2b); 1/521 in the TE DOWN/no Δ mRNA data set (dark green in Figure 2b); 1/107 in TE DOWN/mRNA DOWN data set; 6/152 in the TE UP/mRNA UP data set (pink in Figure 2b); and 6/1,190 in the mRNA UP/no Δ TE data set (orange in Figure 2b). To simplify analysis, these were distributed into three categories ("no change” containing 57/5001 genes, ΔTE containing 40/1304 genes, and ΔmRNA containing 9/1625 genes). The percentage of mTOR-sensitive orthologues within each of these three categories is shown in Figure 2i. Genes that showed changes in both translation and transcription (for example those in the “pink” category) were counted once in each of the above categories. An Exact binomial test was performed in R (binom.test where x was the number of mTOR-sensitive orthologues in a given category, n was the total number of genes in that category and p was the hypothesized probability of mTOR-sensitive genes in the data set to 101/8139) to assess the probability of a given mTOR-sensitive gene belonging to one of these three categories based on the expected probability. In the figure ** indicates p < 0.001, **** indicates p << 0.00001, n.s. indicates p > 0.05 and is not deemed statistically significant.

### Western blot analysis of protein expression

Axolotl limb or mouse digit-tip tissue was harvested as described above and immediately snap-frozen by direct submersion in liquid nitrogen. The samples were powderized with a mortar and pestle under liquid nitrogen or a TissueLyser II chilled to -80°C (Qiagen). The samples were lysed in RIPA buffer for 30 min on ice with intermittent vortexing (0.15M NaCl, 0.05M Tris-HCl pH7.4, 0.005M EDTA pH8, 0.001M EGTA pH8, 0.025M sodium pyrophosphate pH7.4, 0.001M sodium orthovanadate, 0.01M NaF, 0.001M B-glycerol phosphate, 0.1% SDS, 1% NP-40, 0.5% DOC and 1 tablet of cOmplete, Mini, EDTA-free Protease Inhibitor Cocktail (Roche, 11836170001) per 10mL of buffer supplemented with 1X Halt Protease and Phosphatase Inhibitor Cocktail (Thermo Fisher, 78442) prior to use) and cleared by centrifugation at 13,200 rpm at 4°C for 5 min repeated twice. Samples were sonicated with a Bioruptor Sonication System (Diagenode) on medium intensity with a 30 sec ON and 30 sec OFF cycle for 5 min and cleared one more time by centrifugation as described above. The protein concentration of cleared samples was determined with a Pierce BCA Protein Assay Kit (Thermo Fisher Scientific, #23225) with measurements conducted on a GloMax-Multi Plate Reader (Promega, E7081). Equal amounts of protein were diluted in 15 μL or 30 μL total volume with 1X Laemmli SDS-sample buffer (Fisher, 50-196-784), boiled at 95°C for 5 min, chilled on ice and loaded onto 4-20% polyacrylamide gels (Biorad #5671095, #4568096, #4561093, #5671094). The same gel size was used for all replicates of a given experiment. Resolved protein was transferred to a PVDF membrane (Biorad, #170-4273) with the Trans-Blot Turbo Transfer System (Biorad), blocked for 1 h at RT in incubated overnight in primary antibodies against RPS6^Ser235/236^ (Cell Signaling Technology, #4858S), P-RPS6^Ser240/244^ (Cell Signaling Technology, #5364S), P-4EBP1^Thr37/46^ (Cell Signaling Technology, #2855S), RPS6 (Cell Signaling Technology, #2217S), 4EBP1 (Cell Signaling Technology, #9644S), ß-actin (Cell Signaling Technology, #3700S), Rpl19 (Santa Cruz Biotechnology, sc-100830), Rpl7a (Cell Signaling Technology, #2415S), TXN (Aviva, ARP72618), AGR2 (Sigma, AV42290-100UL) and SELENBP1 (Santa Cruz Biotechnology, sc-373726) diluted to 1:1000 in blocking buffer containing 5% bovine serum albumin (Sigma, A9647-100G) and 1xTBST (10mM Tris, 0.15M NaCl, 1% Tween), washed in TBST and incubated with Rabbit (GE Healthcare, GENA934-1ML) or Mouse IgG HRP Linked Whole Ab (GE Healthcare, GENA931-1ML) diluted to 1:10,000 in blocking buffer and incubated at RT for 1 h. Detection was performed on a ChemiDoc MP (Biorad) with Clarity Western ECL Substrate (Biorad, 170-5061) or SuperSignal™ West Femto Maximum Sensitivity Substrate (Thermo Fisher, 34095). In Figures 3, 5 and Extended Data Figures 4-6, the corresponding ß-actin blots are shown directly below blots for target proteins. Because sequential antibody incubation and detection was performed to blot for multiple proteins, of distinct molecular weights, on the same membrane, the ß-actin blots may be shared between multiple blots and therefore are displayed multiple times for clarity. In Figures 3 and 5, lysates were harvested from n=3 axolotls per time point resolved on separate lanes. The dashed line in Figure 3k illustrates the steady state protein expression based on the average (ß-actin normalized) level of a given protein in n=2 DMSO-treated animals at 0 hpa. In Figure 5a and Extended Data Figure 4a, each “mouse” lane represents a distinct pool of digit tips harvested at given time-point.

### INK128 administration studies

To assess the impact of mTOR inhibition on pathway activity (Figure 3f-h), translation (Figure 3i), accumulation of translationally upregulated proteins (Figure 3j-k) and wound closure (Figure 4a-b) and regeneration (Extended Data Figure 3), we applied the same standard protocol whereby axolotls were anesthetized in Tricaine as described above. 100 mM (or 30.933 mg/mL) INK128 (LC Laboratories, I-3344) stock was prepared and stored at -80°C. Axolotls were anesthetized in Tricaine as described above, weighed and 0.025 mg of INK128 per 1g of axolotl body mass, diluted in DMSO (or an equivalent volume of DMSO carrier control) in a total volume of 100 μL (topped up with 0.8x sterile PBS) were administered using an Insulin Syringe with a 28 1/2 G needle (Fisher Scientific, 114-826-205). After 4 h, the animals were anesthetized again and underwent amputation at 24 hpa or 48 hpa for western blot analysis or 12 hpa to assess translation on polysome gradients as described above. For tracking how administration of INK128 prior to amputation impacts the entire process of regeneration, the regenerating limb was briefly imaged under a light microscope at 1, 4, 7, 12, 21, 28, 35 and 42 dpa under anesthesia.

To examine how mTOR inhibition impacts wound closure itself, we immersed white axolotl larvae (AGSC_101L35), 3-5 cm in length, in 10 μM INK128 dissolved in clean tank water (40% Holtfreter’s Salt Solution). After 4 h, the axolotls were anesthetized and amputation was carried out as described above. After amputation, anesthetized larvae were immediately mounted in clean tank water containing anesthetic, and underwent live imaging on a Zeiss Axio Observer Z1 microscope with a Perkin Elmer UltraVIEW Vox spinning disk confocal microscopy system13 until 90 min post-amputation. Representative videos of the process spanning from 30 minutes to 90 minutes post-amputation are included in Extended Data Video 1-2. Percent wound closure for each animal was calculated by taking the difference of percent coverage of skin across the wound site at a starting time point (30 min post-amputation), and at 90 min post-amputation.

To examine whether inhibition of mTOR signaling impacts ROS accumulation after amputation, white axolotl larvae (AGSC_101L35) were immersed in 10 μM INK128, or DMSO control, for 4 h and underwent amputation as described above. The larvae were housed in 10 μM INK128 (or DMSO) containing water and at 36 hpa they were treated with 5 μM 2’,7’-dichlorodihydrofluorescein diacetate (H_2_DCFDA) (Biotium, #10058) for 55 min, and immediately mounted in clean tank water containing anesthetic and imaged. We examined n=6 INK128- treated larvae and n=4 DMSO-treated controls. Single-plane brightfield and H_2_DCFDA fluorescence images of cells lining the wound site of the animal were acquired at a mid-point in the tissue. For quantification, cell boundaries for each image were manually traced in the bright field channel, and then the mean fluorescence intensity of H_2_DCFDA was quantified for each defined cell using ImageJ. The layer of tissue with defined cells is outlined on representative images in Figure 4e.

To assess role of mTOR beyond wound closure (Figure 4h), axolotls underwent the standard amputation protocol as described above. At 4 dpa, wound sites were examined for complete closure on all (12/12) animals and INK128 (or DMSO carrier control) was administered as described above at 0.025 mg of INK128 per 1g under anesthesia. INK128 or DMSO administration were repeated on the same animals at 6, 8 and 10 dpa, injections were administered on alternating sides. Animals were closely monitored and imaged under anesthesia on 4, 14, 28, 42 and 56 dpa. Chronic inhibition of mTORC1 was toxic and only 4/6 animals tolerated INK128 treatment for the duration of the study.

### Multiple Sequence Alignment

To screen for axolotl-specific changes in the mTORC1 pathway, the protein sequences for human mTORC1 pathway components were retrieved from Uniprot and their orthologues were identified in chimp, macaque, mouse, xenopus and zebrafish by protein-to-protein BLAST (https://blast.ncbi.nlm.nih.gov/Blast.cgi). Next, axolotl mTORC1 pathway components were retrieved from https://www.axolotl-omics.org/blast by performing a tblast search against the human sequence. The percentage of amino acid identity between axolotl, human and mouse sequences is reported in Extended Data Table 4. A multiple sequence aligment was performed using Uniprot’s ’Align’ feature using default settings. The alignment was inspected to identify regions with divergent sequences unique to axolotls. This identified insert 1 and insert 2 within axolotl mTOR as regions of interest. To confirm the presence of these inserts across amphibians, which are generally poorly annotated, we searched for mTOR orthologues in the Transcriptome Shotgun Assembly Sequence database. Analysis of the insertion region confirmed that insert 2 was present exclusively within urodele amphibians whereas insert 1 was more broadly conserved across amphibian species. The presence of the insert was further confirmed by examining the sequence reads in our libraries and by amplification by RT-PCR. To conduct a more comprehensive multiple sequence analysis, we retrieved full length mTOR mRNA, and where available protein, sequences for a representative subset of amphibian and non-amphibian species. The mRNA sequences were *in silico* translated with the ExPASy Translate tool and used for multiple sequence alignment using the Uniprot align feature and rendered using the Espript version 3.0 (https://espript.ibcp.fr/ESPript/ESPript/) and the full alignment is shown in Extended Data Figure 6.

### Structure modeling

The predicted model of axolotl mTOR insertion (residues 951-1122) was generated using the transform-restrained Rosetta (trRosetta) algorithm, a deep learning-based protein structure prediction method^83^. All figures were generated using Pymol. A model of the axolotl mTOR insertions was built using the transform-restrained Rosetta (trRosetta) algorithm and superimposed on RHEB-bound mTORC1 dimer (PDB 6BCU). Intriguingly, both insertions reside on mTOR dimerization or effector binding interface, suggesting a potential functional impact on axolotl mTOR. The first insertion (insert 1) in axolotl mTOR contains seven more amino acids (residues 1001-1007) than human mTOR near one of the RHEB switch II region interacting segments. This extended RHEB binding surface in axolotl mTOR might modulate how RHEB allosterically activates the kinase domain. The second insertion (insert 2, residues 1039-1058) in axolotl mTOR is located at the dimerization interface. The predicted model of two insertion 2 suggests a neo-interaction between two axolotl mTOR molecules.

### Generation of chimeric human-axolotl mTOR kinase in HEK293T cells

To model the impact of axolotl specific inserts of the function of mTOR kinase, we used Clustered Regularly Interspaced Short Palindromic Repeats (CRISPR) with CRISPR-associated protein 9 (Cas9) to scarlessly insert these insertions into the human genomic locus encoding mTOR in HEK293T cells which are an established model to study mTOR function^52, 54^. To do this we designed donor templates encoding a human exon 20 carrying an in-frame axolotl insert 1 and a separate donor template encoding a human exon 21 and an in-frame axolotl insert 2. The insertion region was flanked by homology arms ∼300 bp in length (for details see Extended Data Figure 7a). Two guide RNAs (gRNAs) were selected within introns flanking each desired site of insertion and ordered as oligos (IDT). The sequences are listed below. The protospacer adjacent motif (PAM) sequence of each gRNA sites was mutated in the donor templates to prevent re-cutting of the repaired genomic DNA. Note that in all cases the gRNA sequence was intronic. In addition, copies of the gRNA and PAM sequence were appended to the 5’ and 3’ ends of the donor template to enable excision of the donor template from a plasmid as a linear double-stranded DNA “double-cut donor” and increase efficiency of homology-directed repair^84^. A 997 bp donor template containing insert 1 and a 1,363 bp donor template containing insert 2 were synthesized by CODEX DNA (https://codexdna.com/), subcloned into a TOPO cloning vector with a Zero Blunt™ TOPO™ PCR Cloning Kit for Sequencing (Invitrogen, #450031). The gRNAs were subcloned into a PX459 vector (addgene #62988)^85^ and sequenced to confirm correct insertion. HEK293T cells were seeded in a 6-well plate so that they were 70-90% confluent the next day (∼0.8x10^6 - 1x10^6 cells/well in 2 mL of standard growth media containing 10% FBS (Millipore, TMS-013-B) and 1x Penicillin-Streptomycin (Invitrogen, #5140163) in DMEM (Gibco, 11965-118)). Next day, fresh media was added and after 30 min the cells were transfected with CRISPR reagents for insert 2 insertion using the Lipofectamine 3000 Transfection Kit (Invitrogen, L3000001) per manufacturer’s instructions. We used 1600 ng donor template (in TOPO), 400 ng each of 5’ and 3’ gRNA in PX459 vector, 4.8 μL of P3000 reagent and 3.57 μL of Lipofectamine 3000 in a total volume of 250 μL of Opti-MEM(Invitrogen, 31985070). The next day, the cells were split to a 10 cm plate. Puromycin selection was initiated the following day with 1 μg/mL of puromycin dihydrochloride (Sigma P8833-10MG). Puromycin in fresh growth media was added after 24 h. After 48 h of selection, 1/2 of the cells were frozen down, 1/4 were used to extract DNA for PCR verification of insertion within the cell pool and the remaining 1/4 of cells were split to a new 10 cm plate. The following day individual selected cells were trypsinized in 0.05% Tryspin-EDTA for 5min at 37°C, pelleted by centrifugation at 1,000 rpm for 4 min and resuspended in sorting buffer containing HBSS (Gibco, #14175-079), 2% FBS, and 1mM EDTA and filtered. Individual live cells were sorted on into wells of a 96-well plate containing fresh growth media. At least 3 plates were prepared for each construct. After 1 week the growth media was changed once per week and clones reached confluency after two weeks. Clones were screened for insertion by PCR. Homozygous insertion was confirmed by sequencing. After clone containing homozygous insertions of insert 2 were successfully expanded, the above protocol was repeated to introduce homozygous insertion of insert 1. Two independent clones doubly homozygous for insert 1 and insert 2 were generated and used for downstream analysis.

Oligo sequences for gRNAs were as follows for insert 1:

gRNA_ 47163 caccGTCTCAGTAGATAGTGTAGTG & CAGAGTCATCTATCACATCACcaaa, gRNA_47478caccGAGCTGTTACAGTCTTAGTAC & CTCGACAATGTCAGAATCATGcaaa

Oligo sequences for gRNAs were as follows for insert 2:

gRNA_49826 caccGGGGGAGTGAGGAGTTGATCT & CCCCCTCACTCCTCAACTAGAcaaa, gRNA_50114 caccGAAATC AGAAAATCTCTCTGG & CTTTAGTCTTTTAGAGAGACCcaaa

>insert1_donor_template CCTGTACTAAGACTGTAACAGCTTAAGAGACAGGGTCTCACCCTGTCACCCTGCACTCTGTAGAGTGCAGTGGCACGTGCATAGCTCACTACAGCCTTGAACTTCTGGGCTCAAGCATTTCTCCCACCTTAGCATCCAGTAGCTAGAACTACAGGCAAGTGCCCAGCTAATTTTTTGTATTTTTTTTGTAGAGACAGGATCTCGCCATGTTGCCCAGGCTGTTCTTGAAATCTTGGCCTCAAGCAGTCCTCCCGCCTTGGCCTCCCAAAGCACTGAGCCTCCAAACCTAGCCAAGCTTGGGTCTTTGAAATATATTCTCAGTAGATAGTGTAGTGTGCAAACCCATTCCATAGTTGCCTTTATTTGTTCACATGAGCTTAAGGTAAGCCTGGGGGTTCAATGTCCTTCATGATACCAGCTGGTTGACCTCTTGTCCATTTCAGTCCCACCAGCCTTCTCCCCAATTCATGTTCCAACAGCTGGGGATGCTAGTGTCCTTTGTACGGAGTCACATCCGGCCGTACATGGAGGAAATATTCTCTCTCATGAGAGTGAGTAGAAGTTAATGCTTTGGCCTCTTCCATGTTTGGGTCAAGGAAGGCTCAGAAGCAAGTTTGAATGACATAGACTTTTTTTCACGGATCTTTGTAGAGCTGTTACAGTCTTAGTACAGCAAGTGGAACAAAGCCCACTGGATTTTGAGGGGGAGGAAGGGCTGTTGCTCCAGGTTCCCAGGTACAAAACTACAAGGCATGAAGGCTGAAGAGAAATCCTGCAGTAATTGTTCTGCCAGAAATAGACAATTGGGTTATTTGCCTCACACACAACAAAGCAATTTTACTTTAATATCACAAGGGTGCTTTTCTATTTTCATGAAAGCCCTCTTTGTTACTGCTCATAAACCATGAAGGGATTGGGTTTTCTTAGGGCTGTTAAATATGATGGACACGTGTTGGGCACCAAGGAACAAGGTGCCACACTACACTATCTACTGAGA >insert2_donor_template GCGTTTAAACGAATTCGCCCTTAAATCAGAAAATCTCTCTGGAGGGTCTAGAGGGAGGACAGCAAAGCAAAGGAAGTCTTACAACTGCTATCGCTATCAAAGGGAGTGCAAAGGACCCAAGTGGATCATGGAAGGAGAAGAAGAAAGCCTGCATGGGAGAAGTGAGGCTAGAGCCAGGTCTAGAAAGACAGACAGGGCTTAGCTGAGCAAGAAGTGGGATTCTAGGCAGAGGAAGCAGCATGAGCAAAGCCCAGAGGACTGAGGTAGCCTGGTGCATCTGGGGAAATGCAATTGAGTTTGTCCAGTGTAGCTGCAACATAGGACACAAGAAGGGCCCAGTGGGAGGAGGGGAGTGAGGAGTTGATCTCGCCGCAGACCATGCTCACTACAGTTTTGCTTTTCTGGCCATCTTGATTCCTTTGTTCATATATCATCAGAAAGGGACCTGACTCAGCTCCTCTGACTTTTCTCTCTTTGTAGGAATTGAAAACGGATGTCTTGGAGACAACAGATCCATTGAGGACAGACTCAAATAAGGATCACTGGACAACGAATAACCCAATTCAGAGCACGATCATTCTTCTCATTGAGCAAATTGTGGTAGCTCTTGGGGGTGAATTTAAGCTCTACCTGCCCCAGCTGATCCCACACATGCTGCGTGTCTTCATGCATGACAACAGCCCAGGCCGCATTGTCTCTATCAAGGTGAGTAGCCTACGTCATCTTCCAGAGAGATTTTCTGATTTCCTCTGAGTCCCTGGGTGATCAGCTAAAAGCTGAGACCTCATTCTGAGTGACAGGTTGATGCCCATTCCATAAGACAGAATCCCAAGAATACTAATACCCAATGTGTGCAGTTTACAGAATGTCTGTAATCCTCTCTTGATTATCCTTATGTTTTGTATCTGTTTCAATGGATTAATCTTGGGAAATATTTTATCCCAGACTAATTTTCTTTATTTTCCAGCAACGGATTCCTTATCAACTCAAATAAGCACAGAGAAAGCAAAGTAATATGTAAGCAAATAAAATGAGGGGAAGAAAGTGCTATCAAAAGGATATAGTTCAAGGCCATTTAATAAAGAGTTTTCCCAGTCCCCAGAGAACTTTGAATTGTCTACACCACCACCCGCTGCGTGTCCTTAGCCGAGATCAACTCCTCACTCCCCAAGGGCGAATTCGCGGCCGC

### Characterizing the function of chimeric mTOR

To characterize the impact of axolotl inserts on the function of mTOR kinase, we subjected the chimeric HEK293TaxmTOR cells and wildtype controls (both the untreated parental HEK293T line and a line that underwent the CRISPR/Cas9 protocol described above but maintained the wildtype sequence). To assess the nutrient sensitivity of the chimeric mTOR, Wildtype or chimeric HEK293T cells were seeded in 12-well plates at 250,000 to 275,000 cells/mL in standard growth medium (10% FBS, 1x penicillin/streptomycin, DMEM). The next day, the cells were rinsed once in RPMI without amino acids, glucose or serum (US Biological, R9010-01) to remove traces of growth media and starved in RPMI supplemented with 10% dialyzed FBS (Invitrogen, A3382001) and 1x glucose (Invitrogen, A2494001) for 50 min. Next, we prepared a 2-fold dilution series of a 1x “stimulation” solution containing 1x essential amino acids (Invitrogen, 11130051), 1x non-essential amino acids (Invitrogen, 11140050), 1x L-glutamine (Sigma, TMS-002-C), 10% dilayzed FBS and 1x glucose in RPMI. Wildtype or chimeric HEK293T cells were re-fed with 1 mL of “stimulation” solution, ranging in concentration from 1x to 0x, for 30 min. After 30 min, the cells were rinsed with 1x PBS and lysed by incubation with RIPA buffer on ice for 20 min. The cells were then cleared, sonicated and underwent quantitation with BCA and western blotting against phosphorylated and total RPS6 and 4EBP1 as described above in the “Western blotting” section. Three independent titration experiments were performed on the parental HEK293T line and clone chimeric clone C1, a representative blot from one experiment is shown in Figure 5 and Extended Data Figure 7.

To assess the capacity of the chimeric mTOR to sense nutrient starvation and stimulation over time, we seeded cells as described above and starved them for 30 min, 50 min, or 80 min by incubating them in RPMI (US Biological, R9010-01) without amino acids, glucose or serum. Cells were rinsed in 1xPBS and harvested in RIPA buffer at these time-points. After 50 min (the standard starvation interval described in previous work), a subset of starved cells was stimulated by addition of 1x amino acids (essential, non-essential and L-glutamine), 1x glucose and 10% dialyzed FBS. Cells were rinsed with 1xPBS and harvested by lysis in RIPA buffer after 5 min, 10 min, 20 min and 30 min of stimulation. The cells underwent clearing, sonication, BCA quantitation and western blotting against total and phosphorylated RPS6 and 4EBP1 as described above. The data shown represent two independent experiments performed on two independent chimeric and wildtype clones of HEK293T cells. Graphs were plotted in GraphPad Prism 9 and Student’s t-test was used to determine significance.

Lastly, to examine the impact of insertions on lysosomal localization, 200,000 wildtype or chimeric cells were seeded onto fibronectin-coated glass coverslips (Millipore-Sigma, F1141-1MG) in 6-well plates. The next day, each well was gently rinsed once with RPMI and then starved for 50 min in 2 mL of RPMI (without amino acids, glucose or FBS) with 20 μM Lysotracker Red DND-99 (Invitrogen, L7528). After 50 min of starvation, the cells were stimulated by addition of 1x essential amino acids, 1x non-essential amino acids, 1x L-glutamine, 1x glucose, and 10% dialyzed FBS directly to the medium and incubated for 10 min. For control cells, the Lysotracker was added directly to the existing media. The cells were then rinsed with 2 mL of 1xPBS containing Lysotracker and fixed in 4% paraformaldehyde in PBS for 15 min at RT. After incubation the cells were rinsed twice with 1xPBS and permeabilized for 10 min at RT with 0.1% saponin. They were rinsed with 1x PBS again and incubated with 50 μL of blocking solution (1x PBS, 0.5% goat serum) with 1:300 anti-rabbit mTOR (Cell Signaling Technology, #2983S) for 1 h at RT. The cells were washed three times with 1xPBS and incubated with 1:1000 Alexa 488 secondary antibody (Invitrogen, A11070) and 1:1000 DAPI for 45 min at RT in the dark. The cells were rinsed three times in 1xPBS and mounted on slides using Fluoromount-G Slide Mounting Medium (Southern Biotech, 0100-01). Slides were images on a Zeiss Axio Observer Z1 microscope coupled to the Perkin Elmer UltraVIEW Vox spinning disk confocal microscopy system. Analysis was performed using custom code written in Python 3.7. The mean fluorescence intensity of mTOR (488 nm laser channel) was measured within lysosomes defined by Lysotracker (568 nm laser channel) using local adaptive thresholding in individual cells defined by manually outlined cell boundaries. Cell boundaries were carefully drawn in Fiji/ImageJ based on DAP (405 laser channel) and mTOR staining. For each cell line and condition, the lysosomal intensity is expressed relative to the mean steady state intensity for that specific cell line. Significance assessed with one-way ANOVA calculated in GraphPad Prism 9.

### Cell Viability Assay

Wildtype HEK293T cells or HEK293T^axmTOR^ sells were seeded at a density of 100,000 cells/mL on 96-well plates, allowed to attach at least 12 h and treated with INK128 (Selleckchem, #S2811) at indicated concentrations for 48 h in standard growth media. A CellTiter-Glo Cell Viability Assay (Promega, G9241) was performed following the manufacturer’s instructions and absorbance was measured at 490nm using a GloMax-Multi Plate Reader (Promega, E6521). Three independent experiments were performed. The IC50 values for each replicate was calculated in Prism 9 by performing nonlinear regression for inhibitor vs response (variable slope-four parameters). The average of three IC50 values +/- standard deviation is shown in Figure 5i. Significance was calculated using Student’s t-test.

### Data availability

Raw data supporting this work will be deposited to Gene Expression Omnibus prior to publication.

### Computer code

All custom code is available upon request from the corresponding author.

## Acknowledgements

The authors would like to thank members of the Barna laboratory for critical feedback and discussion of this work; S. Arulmani and S. Stern for preliminary contributions; A. Valdefiera, S. Ahmadi, S. Jensen, and D. Chu of the Stanford Veterinary Services Center for animal husbandry and veterinary support; V. Natu and J. Coller of the Stanford Functional Genomics Facility for sequencing; A. Chekholko of the Stanford Genomics Center for IT system support; S. Floor (UC Berkley) for protocol assistance; R. Voss of the Ambystoma Genetic Stock Center (University of Kentucky) for training and materials. Funding: this work was supported by New York Stem Cell Foundation grant NYSCF-R-I36 (M.B.), New York Stem Cell Robertson Investigator (M.B.), NIH grant 1R01HD086634 (M.B.), Stanford Discovery Innovation Fund in Basic Biomedical Sciences (M.B.), Canadian Institutes of Health Research postdoctoral fellowship (O.Z.), K99/R00 Pathway to Independence Award NICHD grant 5K99HD099787-02 (O.Z.), National Institutes of Health Developmental Biology training grant 5T32GM007790-41 (H.D.R.). O.Z. is a Simon’s fellow of the Helen Hay Whitney Foundation. M.B. is a NYSCF Robertson Investigator.

## Author contributions

O.Z. and M.B. conceived the project and designed experiments; M.B. supervised the project; O.Z. performed amputation, sucrose gradient fractionation, polysome sequencing and data analysis, western blot analysis, INK128 incorporation studies, generation of chimeric cell lines by CRISPR/Cas9, AA titration, immunofluorescent staining on cells and tissues, OPP incorporation studies and training and supervision for H.D.R. and L.S.; H.D.R. designed and performed ROS detection and INK128-incorporation and live imaging studies, carried out mTOR imaging and data analysis and acquisition in cells and tissues; L.S. optimized mTOR imaging and generated preliminary data on lysosomal localization of mTOR; S.D. performed sequence analysis and structure modelling; Z.Z. generated a pipeline for semi-automated mTOR localization analysis; D.K.-O. performed cell viability assays; D.R. supervised D.K.-O. and provided critical feedback on experimental design; K.M.S. supervised S.D. and provided critical feedback and assistance with experimental design. O.Z. and M.B. wrote the manuscript with input from all the authors.

## Extended Data Figure legends

**Extended Data Figure 1:**
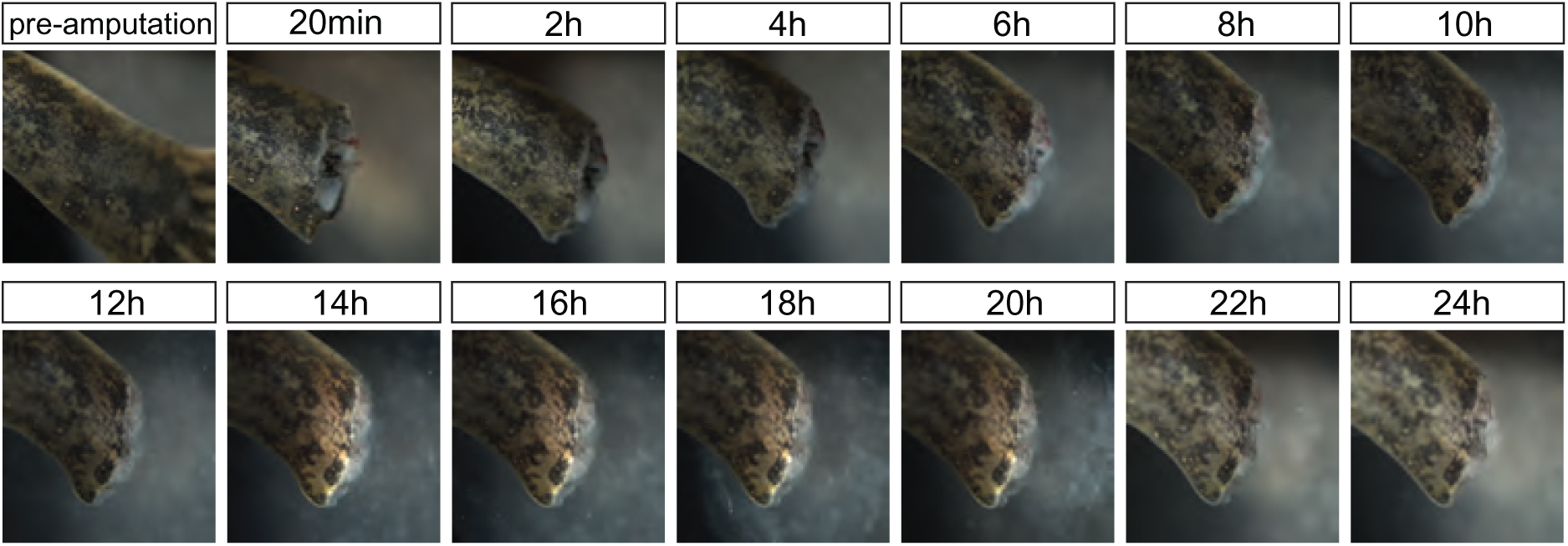
Rapid epithelial migration drives wound closure within 24 h of limb amputation in axolotls. Snapshots of wound closure in a wildtype sub-adult axolotl imaged over the course of 24 h.

**Extended Data Figure 2:**
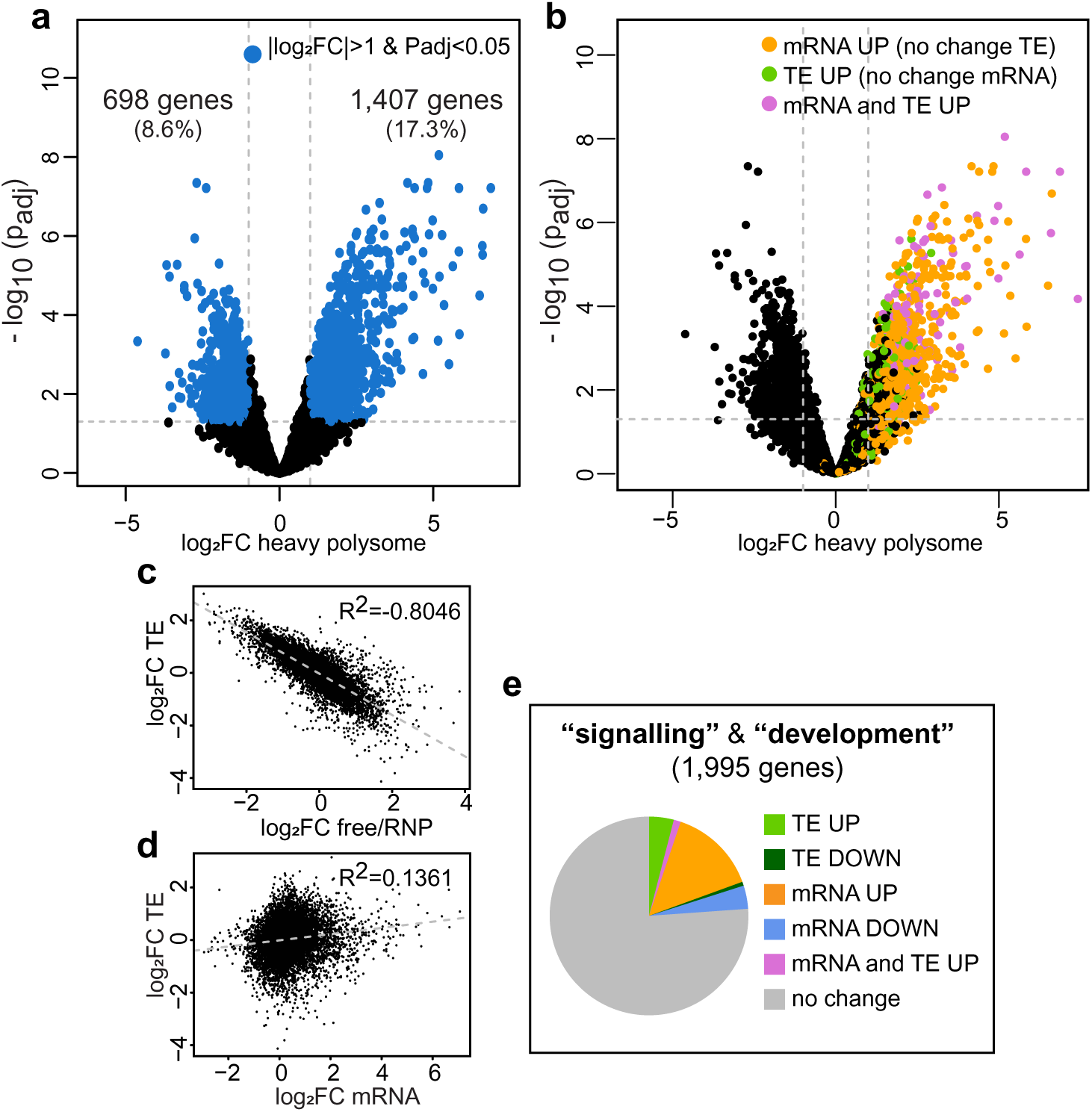
Polysome sequencing identifies translationally regulated mRNAs. **a**, Scatterplot of 8,139 transcripts in our data set. The x-axis shows (log_2_) fold change (FC) in the heavy polysome fractions between 0 h and 24 h post-amputation. Y-axis shows (-log_10_) adjusted p-value (p_adj_). Blue dots indicate transcripts with p_adj_ < 0.05 and two-fold change in the heavy polysome. **b**, The scatterplot from panel (a) colored to emphasize that transcripts with increased reads in heavy polysome include transcripts that are upregulated at the level of transcription (orange), translation (green) or both (pink) as defined in Figure 2b, therefore change in heavy polysome on its own does not adequately describe the provenance of actively translated transcripts. **c**, Strong negative correlation between (log_2_) FC in TE on y-axis and (log_2_) FC in the free/RNP fraction. **d**, Poor correlation between (log_2_) FC in TE on y-axis and (log_2_) FC in mRNA abundance between 0 h and 24 h shown on x-axis. **e**, Distribution of 1,995 genes with annotated roles in “signaling” and “development” across expression categories defined in Figure 2b.

**Extended Data Figure 3:**
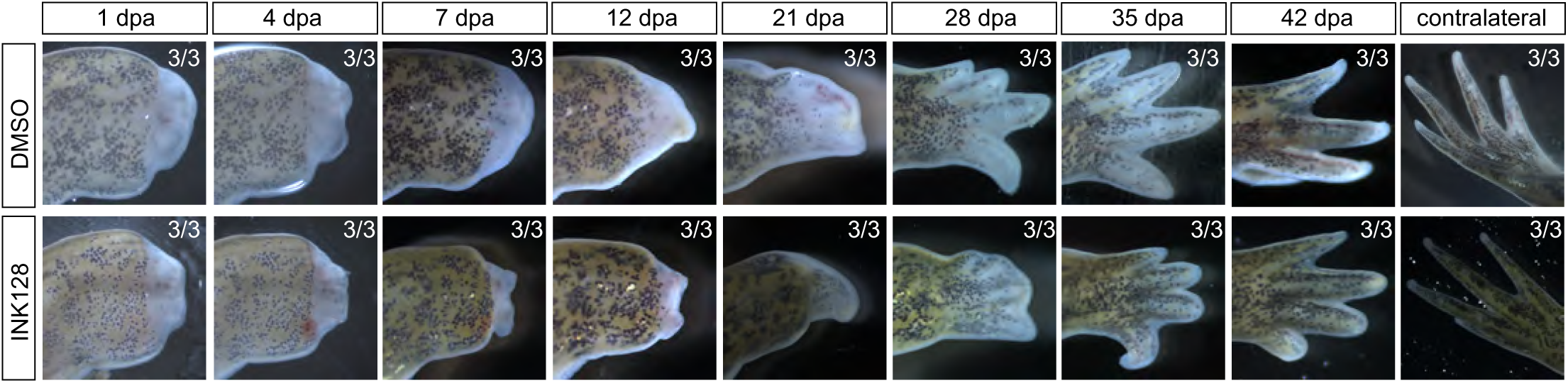
INK128 treatment inhibits regeneration. Representative images of limb regeneration in axolotls treated with DMSO or INK128 4h before amputation. The INK128-treated animals tracked in this analysis all belong to the 37.5% of animals with partial wound closure at 24h post-amputation referred to in Figure 4b.

**Extended Data Figure 4:**
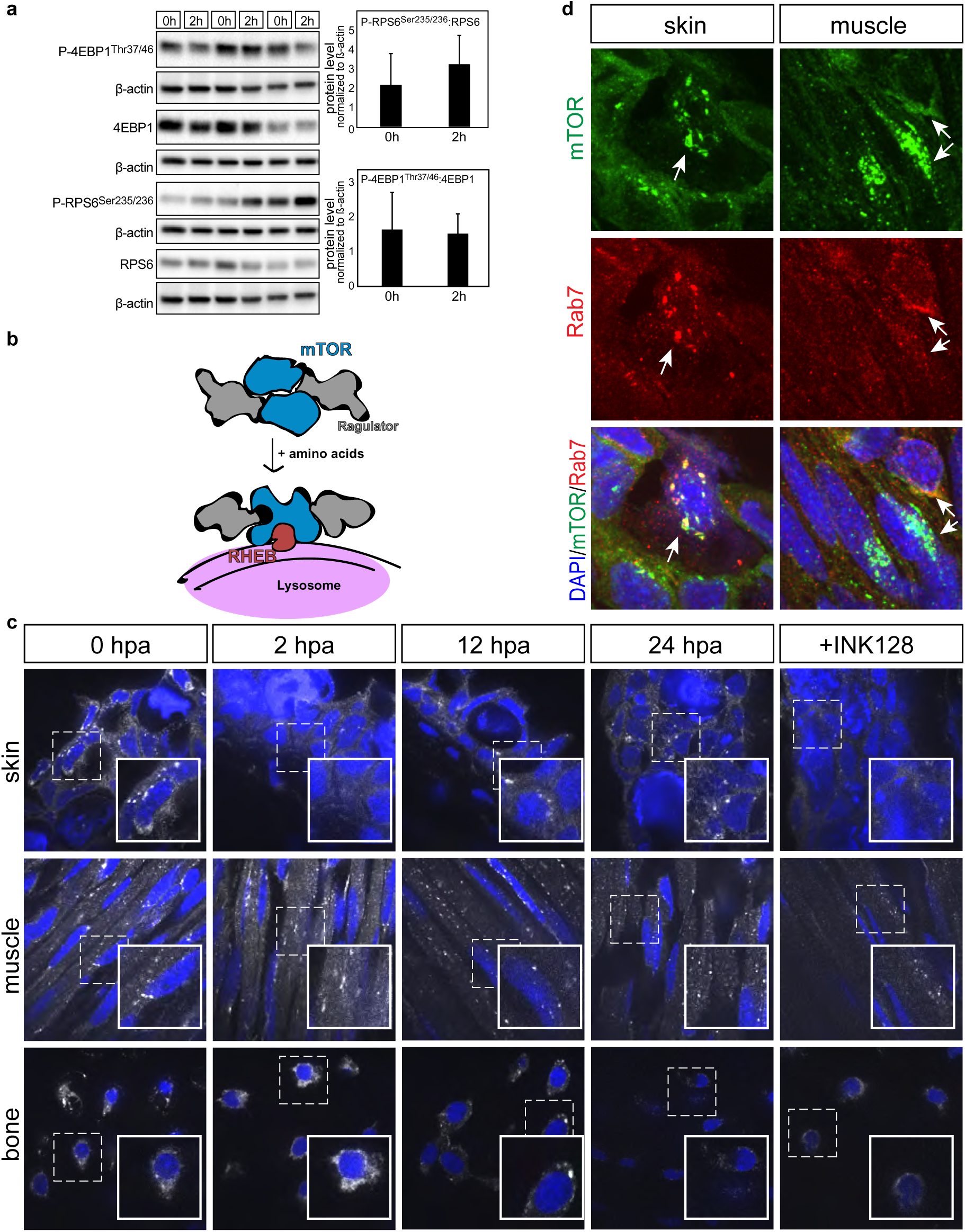
Altered mTOR activity in axolotls and mice. **a**, Western blot depicts mTOR activity measured as phosphorylation of P-4EBP1^Thr37/46^ and P-RPS6^Ser235/236^ in mice at 0h and 2h post-amputation. Each lane represents a distinct pool of digit tips harvested at given time-point. Quantitation shows ratio of phosphorylated to total protein normalized to ß-actin. P-4EBP1^Thr37/46^ and RPS6 were sequentially blotted on the same membrane and share ß-actin blot. 4EBP1 and P-RPS6^Ser235/236^ were sequentially blotted on another membrane and share ß-actin. **b**, Schematic depicts amino acid dependent translocation of mTORC1 to the lysosome is required for pathway activation. **c**, Immunofluorescence staining of axolotl tissues depicts mTOR localization (white/grey) and nuclei (DAPI/blue) in cells of the skin, muscle and bone near the wound site at 0, 2, 12, and 24h post-amputation (hpa). The punctate mTOR localization is lost upon treatment with INK128. **d**, Rab7 (red), mTOR (green) and nuclear stain (DAPI/blue) in axolotl tissues at 24h post-amputation illustrate co-localization of mTOR to lysosomes.

**Extended Data Figure 5:**
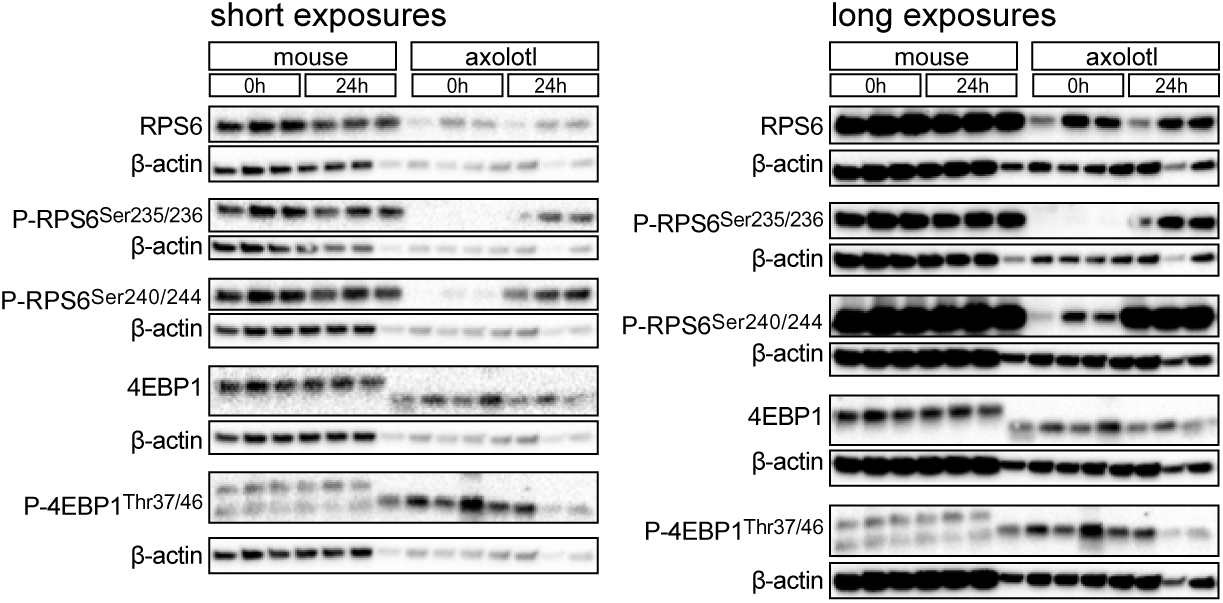
Protein expression in axolotls and mice. Short (∼1ms) and long exposures are provided for the blots shown in Figure 5a to illustrate that differences in mTOR responsiveness to amputation are specific to axolotls independent of exposure time.

**Extended Data Figure 6:**
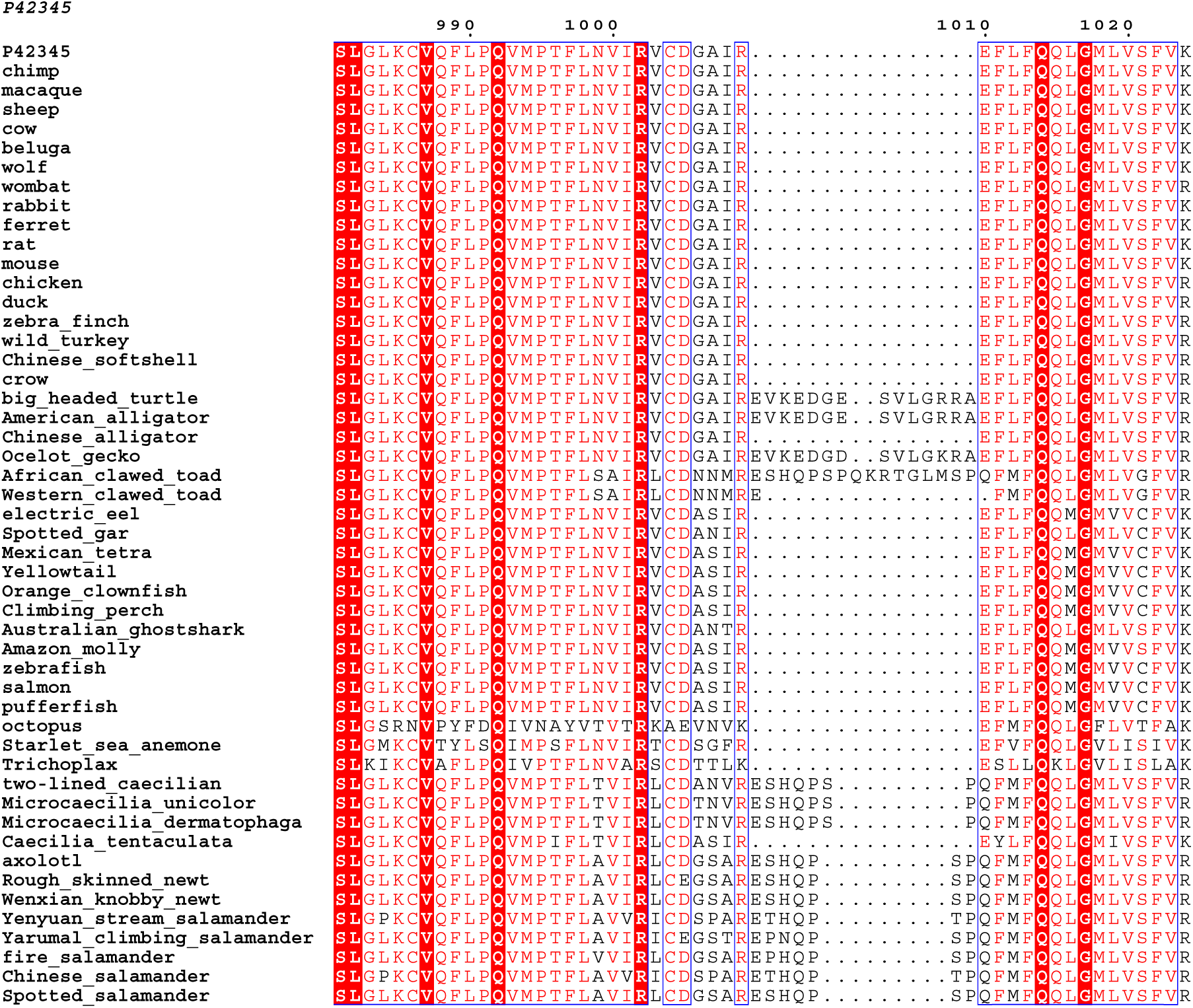
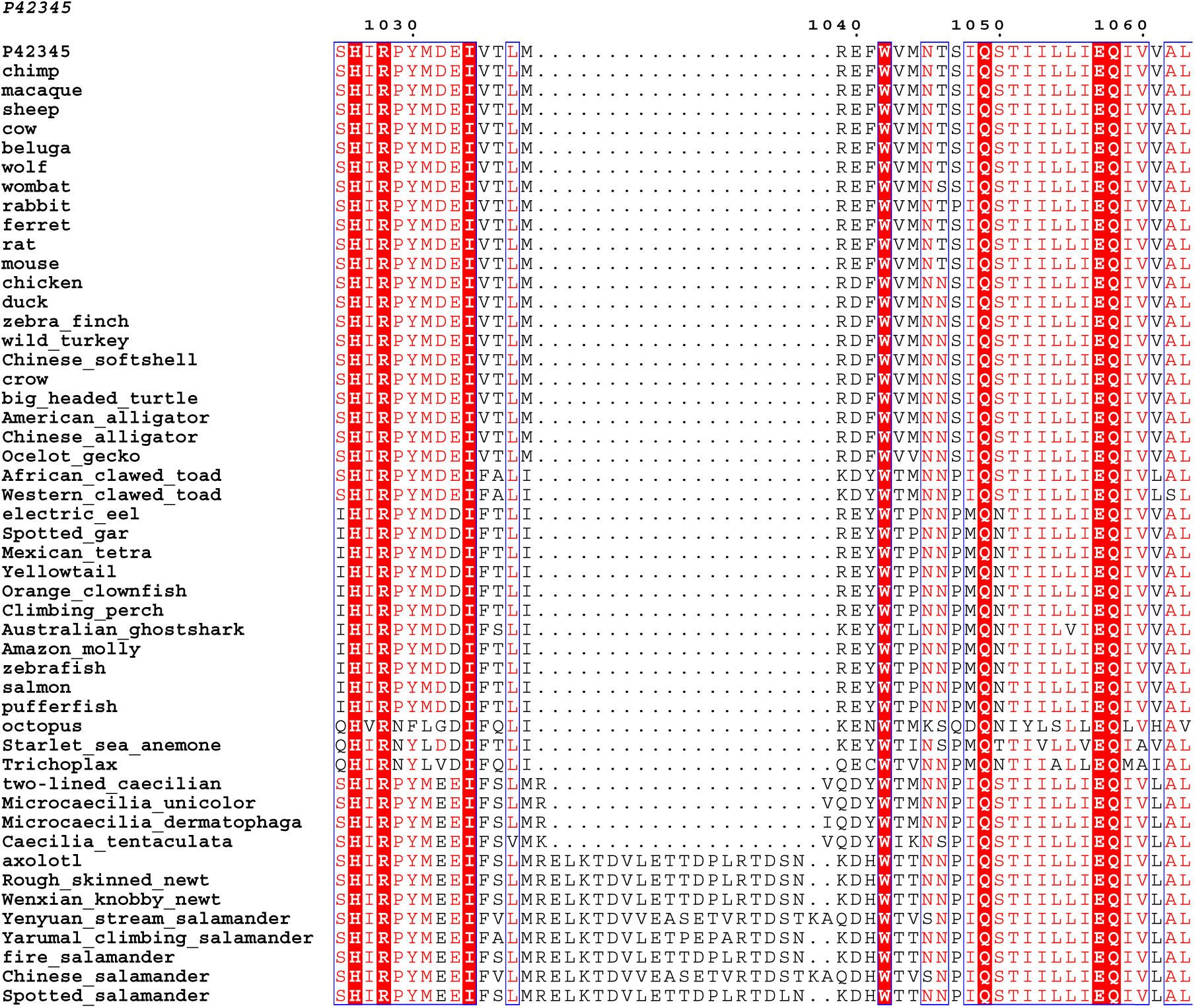
Multiple Sequence Alignment of metazoan mTOR. An alignment of predicted full-length mTOR sequences from 50 metazoan species conducted using Clustal Omega.

**Extended Data Figure 7:**
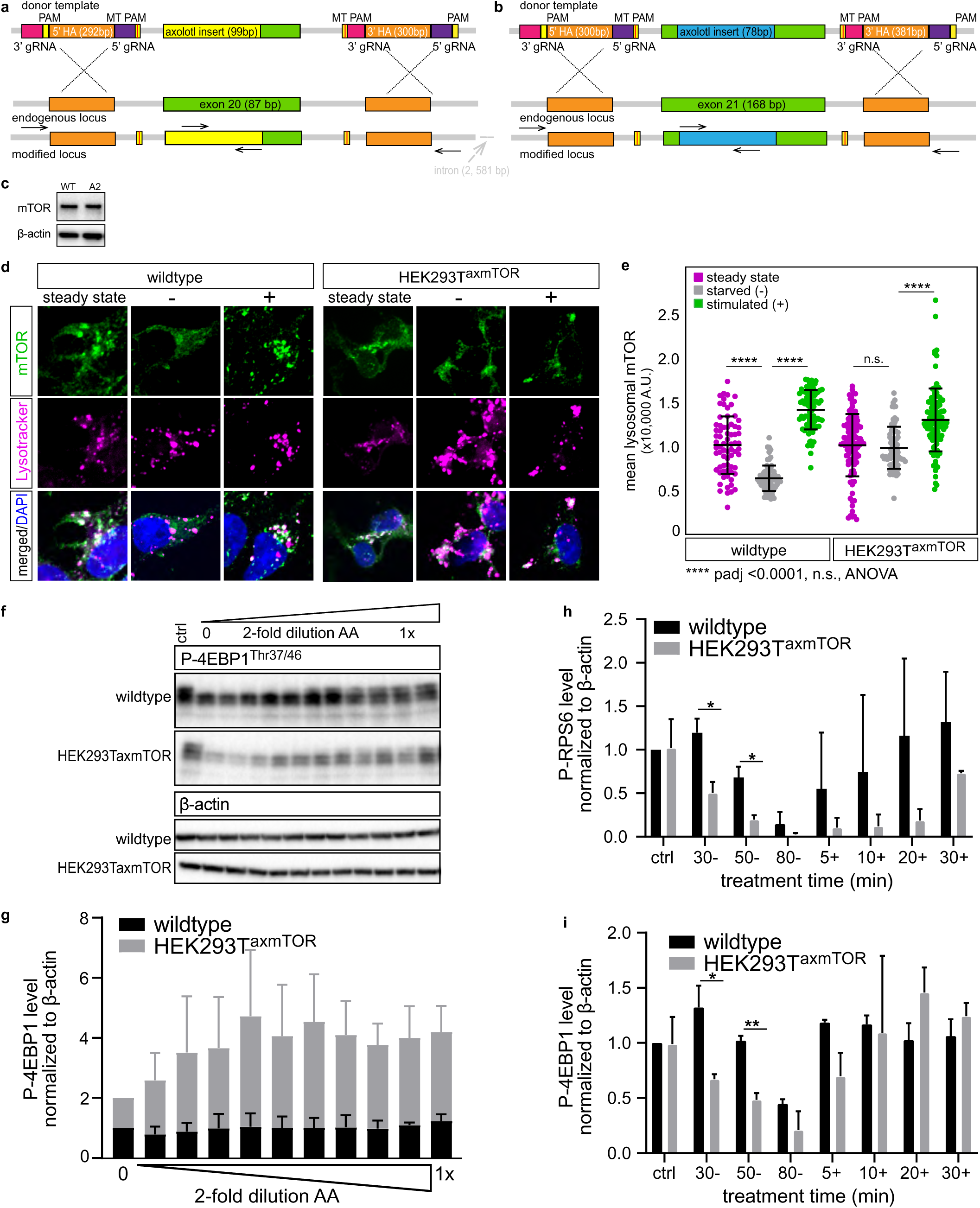
Axolotl insertions promote lysosomal retention of mTOR. **a**, Schematic depicts/Cas9 targeting strategy to introduce axolotl insert 1 in-frame into exon 20 of human mTOR in HEK 293T cells. gRNA (guide RNA site), HA (homology arm), PAM (protospacer adjacent motif) MT PAM (mutated protospacer motif). **b**, Schematic depicts CRISPR/Cas9 strategy to introduce axolotl insert 2 in-frame into exon 21 of human mTOR in HEK 293T cells. **c**, Western blot analysis illustrates steady-state expression of mTOR protein at steady state in wildtype HEK 293T cells (WT) and HEK 239T^axmTOR^ (A2) clone. **d**, Immunofluorescence staining of mTOR (green), lysosomes (Lysotracker in magenta) and nuclei (DAPI) in wildtype and HEK 239T^axmTOR^ cells at steady state, upon starvation (-), and stimulation. **e**, Quantitation of mean mTOR intensity within lysosomes. For each cell line and condition, the lysosomal intensity is expressed relative to the mean steady state intensity for that specific cell line. Significance assessed with one-way ANOVA. **f**, Representative western blot (n=3) of amino acid (AA) titration experiment illustrates greater sensitivity of P-4EBP1^Thr37/46^ phosphorylation in HEK 293T^axmTOR^ to amino acid concentration. **g**, Quantitation P-4EBP1^Thr37/46^ level normalized to ß-actin. **h**, Quantitation of P-RPS6^Ser235/236^ level and **i**, P-4EBP1^Thr37/46^ level each normalized to ß-actin in wildtype and HEK 239T^axmTOR^ in wildtype and HEK 239T^axmTOR^ cells in 2 independent experiments. Significance of response to starvation assessed with Student’s t-test, * is p<0.05, ** is p<0.01.

